# Disassembly of the *Escherichia coli* AcrABZ-TolC efflux pump by ligand-mediated disruption of TolC-AcrA interfacial contacts

**DOI:** 10.1101/2025.02.18.638599

**Authors:** Tania Szal, Emmanouela Petsolari, Janis Veliks, Cristina D. Cruz, Laura Paunina, Marina Madre, Fatima-Zahra Rachad, Philipp Lewe, Susanne Witt, Päivi Tammela, Aigars Jirgensons, Ben F. Luisi, Björn Windshügel

## Abstract

The outer membrane factor TolC is an essential component of various efflux pump complexes in *E. coli* and represents a potential target for antibiotic adjuvants. By means of a virtual screen for TolC-binding compounds, we identified the kinase inhibitor CEP- 37440 to shift the minimum inhibitory concentration of antibiotics piperacillin and levofloxacin in *E. coli*. To determine the substructure relevant for TolC binding, a hit deconstruction approach was applied, resulting in a fragment-like compound with low affinity for TolC and AcrB (LP-115). Dynamic light scattering revealed LP-115 to reduce the hydrodynamic radius of AcrABZ-TolC, indicating a disassembly of the efflux pump complex. A cryo-EM structure demonstrated LP-115 to bind at the TolC-AcrA interface within the AcrABZ-TolC complex, thereby disordering the interface and inducing a closed conformation of TolC. Our results suggest that ligand-mediated TolC-AcrA interface disruption represents a novel mechanism of efflux pump inhibition.

## Introduction

In Gram-negative bacteria, tripartite efflux pumps extrude potentially toxic compounds from the periplasmic space and the cytosol^1^. The substrate spectrum of these complexes is diverse and includes secondary metabolites, siderophores as well as various classes of antibiotics^2,3^. The extrusion of antibiotics by efflux pumps results in sub-inhibitory drug concentrations in the bacterial cell and eventually impairs therapeutic efficacy. Therefore, efflux pumps are considered as a major factor of antimicrobial resistance^4^. The composition of the tripartite efflux pump complex includes a transporter protein, which is embedded in the inner membrane and involves members of the resistance-nodulation-cell division (RND), the ATP-binding cassette (ABC), or the major facilitator superfamilies (MFS), a periplasmic adapter protein, as well as an outer membrane factor (OMF)^5^.

One of the most studied efflux pumps is the tripartite complex AcrAB-TolC in *E. coli*, which has been shown to extrude, among others, rifampicin, erythromycin, β-lactams, and tetracycline^3^. X-ray crystallography and cryogenic electron microscopy (cryo-EM) have provided detailed insights into the structure of all AcrAB-TolC components as well as their assembly^6,7^. The OMF TolC is a homotrimer that is composed of a beta-barrel embedded in the outer membrane and an alpha-barrel that protrudes approximately 100 nm into the periplasmic space (Koronakis 2000)^6^. TolC is a component of several efflux pump systems. In *E. coli*, up to thirteen different tripartite efflux pumps involving RND, ABC, or MFS transporter proteins utilize TolC^1^.

Efflux pump inhibition represents an attractive approach for restoring the efficacy of antibiotics. In recent years, several small molecules have been reported to inhibit AcrB, thereby shifting the minimum inhibitory concentration (MIC) of various antibiotics^8,9^. In addition, several compounds have been demonstrated to bind to AcrA and to have a synergistic effect in combination with antibiotics^10,11^. Recently, several colicin E1 fragments have been shown to interact with TolC and to improve the susceptibility of *E. coli* to several antibiotics^12^. Despite the functional importance of TolC for all known efflux pumps in *E. coli*, so far, no organic small molecule has been reported to specifically bind to the OMF TolC. A few inorganic cations such as hexaamminecobalt (HC) have been demonstrated to bind to and block TolC^13^. The X-ray crystal structure of TolC in complex with HC revealed that the compound binds close to the periplasmic tip in the channel lumen and forms several salt bridges with Asp374^13^. However, the substance itself does not shift the MIC of the antibiotic erythromycin or fusidic acid^14^.

As an alternative to the conventional drug discovery approach, drug repurposing has gained increasing interest because it may reduce development costs and shorten the time to approval^15^. Trimethoprim and epinephrine have been shown to have a synergistic effect in combination with several antibiotics against a set of Gram-negative bacteria and to inhibit the efflux of the fluorescent dye Hoechst 33342^16^. More recently, several approved drugs have been reported to inhibit AcrB in *E. coli* and *Salmonella enterica* serovar Typhimurium^17^.

In the present study, we aimed to identify TolC-targeting small molecules that improve the efficacy of different antibiotics in *E. coli*. Based on an *in silico* repurposing screen, we predicted a kinase inhibitor, CEP-37440, to bind favorably to the periplasmic tip of TolC. Experimental validation revealed the compound to shift the MIC of piperacillin and levofloxacin. Moreover, microscale thermophoresis measurements showed binding to AcrB and TolC. To identify the structural motif responsible for the interaction, a hit deconstruction approach was chosen, by which fragments of the CEP-37440 were tested for TolC binding. This led to the identification of a low molecular weight compound that revealed a weak interaction with TolC and AcrB. Dynamic light scattering revealed LP- 115 to destabilize the efflux pump complex. Structural studies of AcrABZ-TolC using cryo-EM revealed LP-115 to bind at the TolC-AcrA interface and to promote structural rearrangements compared to the unliganded complex. Our results indicate that the TolC-AcrA interface can be specifically addressed by small molecules, thereby expanding the druggable components of tripartite efflux pumps to the outer membrane factors.

## Results

### Virtual screening for TolC-binding compounds

To identify small molecules targeting TolC, a structure-based virtual screening campaign was conducted using the X-ray crystal structures of ligand-free and hexaamminecobalt- bound TolC. Initially, the Site Finder method was utilized to identify and to prioritize potential ligand-binding sites. The calculation suggested the propeller-shaped cavity at the periplasmic tip of TolC, as well as an area close to the hexaamminecobalt-binding site, as suited for small molecule binding. Several areas with a propensity for a ligand binding score above 1 indicate the potential to harbour a small molecule that may either block the channel or interfere with AcrA binding (Fig. S1).

To select compounds with an increased chance of outer membrane penetration, a drug repurposing library containing approximately 6,500 approved and investigational drugs was searched for compounds carrying a positively charged nitrogen atom. The selected compounds were docked into the periplasmic tip of TolC. Based on the analysis of the docking scores and visual inspection of the binding modes of the top-ranked docking poses, eight compounds were selected for experimental validation (Table S1).

### Experimental validation of virtual hits

Initially, the virtual hits were evaluated with respect to their potential to functionally interfere with the efflux pump by increasing the accumulation of the AcrAB-TolC substrate and fluorescent dye H33342 in *E. coli* BW25113 (Table 1). Initial tests were performed at a compound concentration of 100 µM and with phenylalanine arginine beta-naphthylamide (PAβN) as a positive control. PAβN at 100 µg/mL has been shown to reduce the MIC of ciprofloxacin and levofloxacin in *E. coli*^18,19^. All compounds at least partially blocked H33342 efflux from *E. coli* BW25113. Zolmitriptan and xamoterol displayed the least interference with H33342 efflux.

**Table 1.**
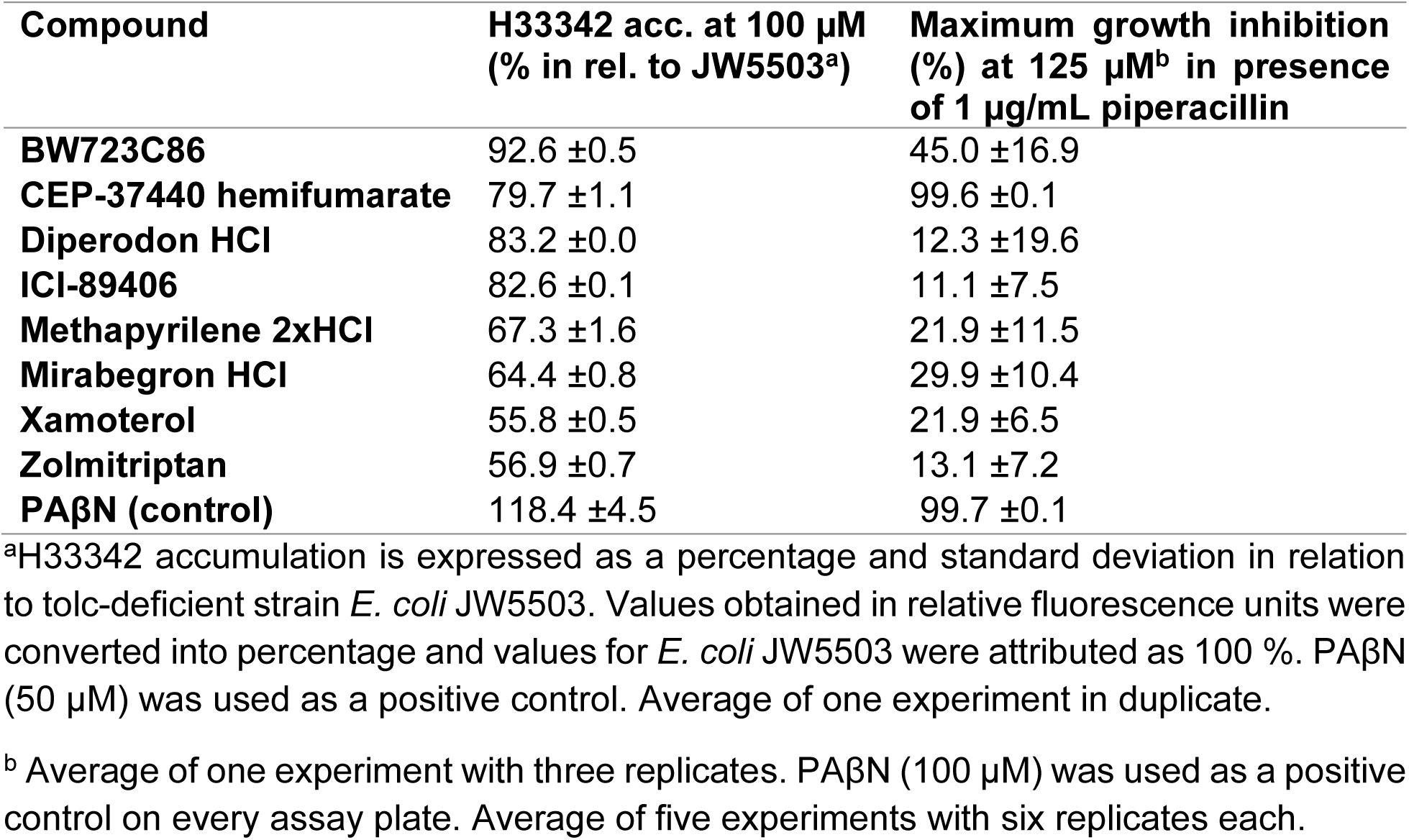
Accumulation of fluorescent dye Hoechst 33342 in *E. coli* BW25113 and inhibition of *E. coli* BW25113 growth in combination with piperacillin (± standard deviation)

| Compound | H33342 acc. at 100 μM<br>(% in rel. to JW5503<sup>a</sup>) | Maximum growth inhibition<br>(%) at 125 μM<sup>b</sup> in presence<br>of 1 μg/mL piperacillin |
| --- | --- | --- |
| **BW723C86** | 92.6 ±0.5 | 45.0 ±16.9 |
| **CEP-37440 hemifumarate** | 79.7 ±1.1 | 99.6 ±0.1 |
| **Diperodon HCl** | 83.2 ±0.0 | 12.3 ±19.6 |
| **ICI-89406** | 82.6 ±0.1 | 11.1 ±7.5 |
| **Methapyrilene 2xHCl** | 67.3 ±1.6 | 21.9 ±11.5 |
| **Mirabegron HCl** | 64.4 ±0.8 | 29.9 ±10.4 |
| **Xamoterol** | 55.8 ±0.5 | 21.9 ±6.5 |
| **Zolmitriptan** | 56.9 ±0.7 | 13.1 ±7.2 |
| **PAβN (control)** | 118.4 ±4.5 | 99.7 ±0.1 |

<sup>a</sup>H33342 accumulation is expressed as a percentage and standard deviation in relation to tolC-deficient strain *E. coli* JW5503. Values obtained in relative fluorescence units were converted into percentage and values for *E. coli* JW5503 were attributed as 100 %. PAβN (50 μM) was used as a positive control. Average of one experiment in duplicate.
<sup>b</sup> Average of one experiment with three replicates. PAβN (100 μM) was used as a positive control on every assay plate. Average of five experiments with six replicates each.

In a second step, the compounds were tested for potential synergistic activity with the antibiotic piperacillin at a sub-MIC of 1 µg/mL in *E. coli* BW25113 (Table 1). Only the kinase inhibitor CEP-37440 demonstrated a synergistic effect with piperacillin at a concentration of 125 µM. PAβN showed the same synergism at 100 µM. Further dose-response experiments with CEP-37440 revealed the compound to act synergistically with piperacillin activity at 50 µM. Consequently, the minimum potentiating concentration required to decrease the MIC of piperacillin by 4-fold (MPC_4_) was 50 µM (Table 2). In combination with levofloxacin, the MPC_4_ of CEP-37440 was 100 µM. To assess the antibacterial activity of CEP-37440 itself, the MIC was determined. CEP-37440 did not inhibit bacterial growth of *E. coli* BW25113 and the *tolc*-deficient strain JW5503 up to a concentration of 125 µM. Furthermore, the specificity of CEP-37440 on the modulation of the *E. coli* efflux pump was investigated by performing the checkerboard assay using the *tolc*-deficient *E. coli* strain JW5503. In contrast to *E. coli* BW25113, no synergistic effect with piperacillin occurred. PAꞵN demonstrated a synergistic effect, resulting in a MPC_4_ of 25 µM which is already known to be related to outer membrane permeabilization^20^.

**Table 2.** Inhibition of *E. coli* BW25113 and JW5503 growth by compounds in absence and presence of piperacillin or levofloxacin.

|  | *E. coli* BW25113 |  |  |  |  | *E. coli* JW5503 |  |  |
| --- | --- | --- | --- | --- | --- | --- | --- | --- |
|  | Piperacillin |  |  | Levofloxacin |  | Piperacillin |  |  |
| Compound | MIC<sup>a</sup> | MPC<sub>4</sub><sup>b</sup> | MIC/<br>MPC<sub>4</sub> | MPC<sub>4</sub> | MIC/<br>MPC<sub>4</sub> | MIC | MPC<sub>4</sub> | MIC/<br>MPC<sub>4</sub> |
| CEP-37440 | >125 µM | 50 µM | >2.5 | 100 µM | >1.3 | >75 µM | >75 µM | - |
| PAβN | >200 µM | 100 µM | >2 | 100 µM | >2 | 100 µM | 25 µM | 4 |

<sup>a</sup>MIC: minimum inhibitory concentration was defined as the of compound concentration that inhibited *E. coli* growth by ≥90 %. If MIC was not achieved, the value is represented by the highest compound concentration tested. Each value represents the average of four experiments performed with three replicates each.
<sup>b</sup>MPC<sub>4</sub>: minimum potentiating concentration was defined as the concentration of compound that decreased the MIC of antibiotics by 4-fold (i.e. piperacillin from 4 µg/mL to 1 µg/mL and from 0.25 µg/mL to 0.0625 µg/mL in *E. coli* BW25113 and JW5503, respectively; and levofloxacin from 0.04 µg/mL to 0.01 µg/mL in *E. coli* BW25113). Each value represents the average of two experiments performed with one to three replicates each.
-: no potentiation of piperacillin observed

### Binding of CEP-37440 to efflux pump proteins

To investigate the interaction of CEP-37440 with the efflux pump AcrAB-TolC, we assessed its binding to each efflux pump subunit using microscale thermophoresis (MST). CEP-37440 bound to TolC and AcrB with high micromolar affinity (Figs. 1a and 1b), whereas the compound did not bind to AcrA or tobacco etch virus protease (TEVp), a sequentially unrelated protein (Fig. 1c).

**Fig. 1.**
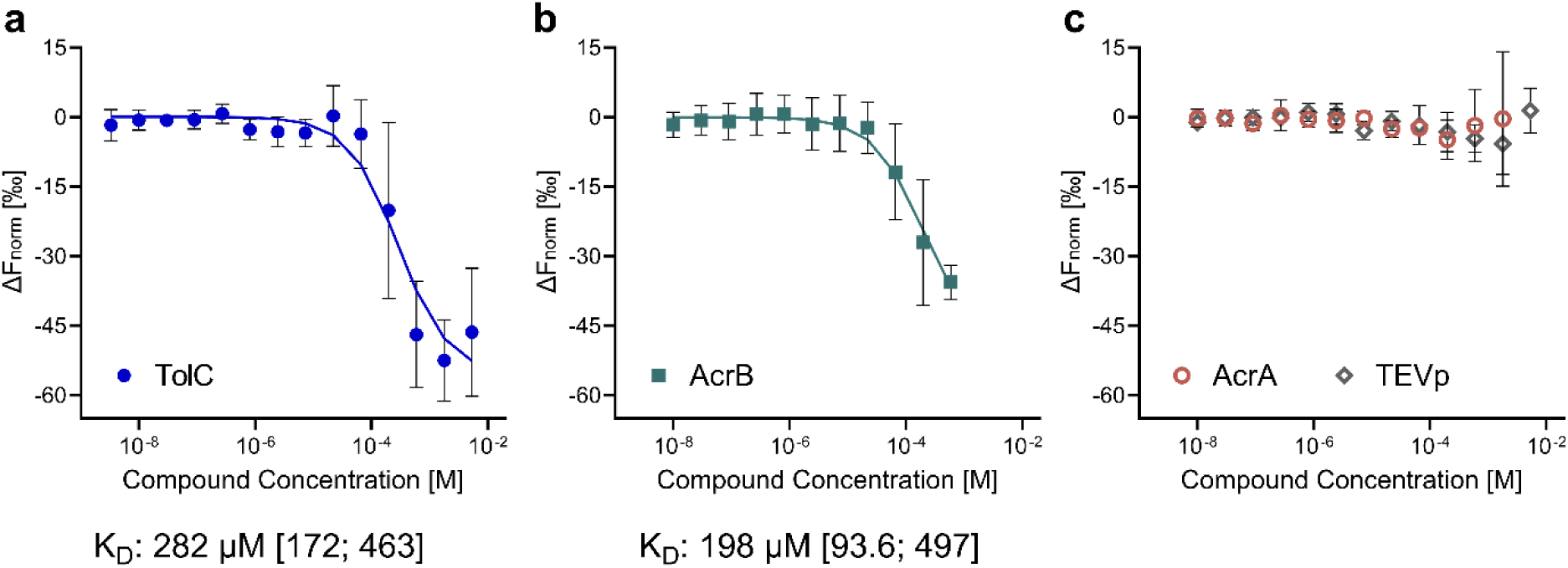
Protein-ligand interaction studies for CEP-37440 and the individual efflux pump subunits. **a** MST signals of 5 µM TolC in combination with a 1:3-dilution series of CEP-37440 (5.3 mM - 3.3 nM). **b** MST signals of 1.25 µM AcrB and **c** 5 µM AcrA and TEVp in combination with a dilution series of CEP-37440 (5.3 mM - 10 nM; for AcrB, high concentrations had to be excluded because of quenching). Each data point is represented as the mean ± SD (n=5). The K_D_ for TolC and AcrB is specified in the graph with its respective asymmetric 95 % confidence interval. Data is derived from MST Traces shown in Fig. S2.

### CEP-37440 deconstruction into fragment-like compounds

Owing to the modest synergistic effect of CEP-37440, as well as its large size (580.1 Da), the potential for optimization is limited. Therefore, we systematically deconstructed CEP- 37440 into fragment-like substructures to identify the functional groups responsible for TolC binding. Altogether five CEP-37440 fragments were synthesized (Fig. 2).

**Fig. 2.**
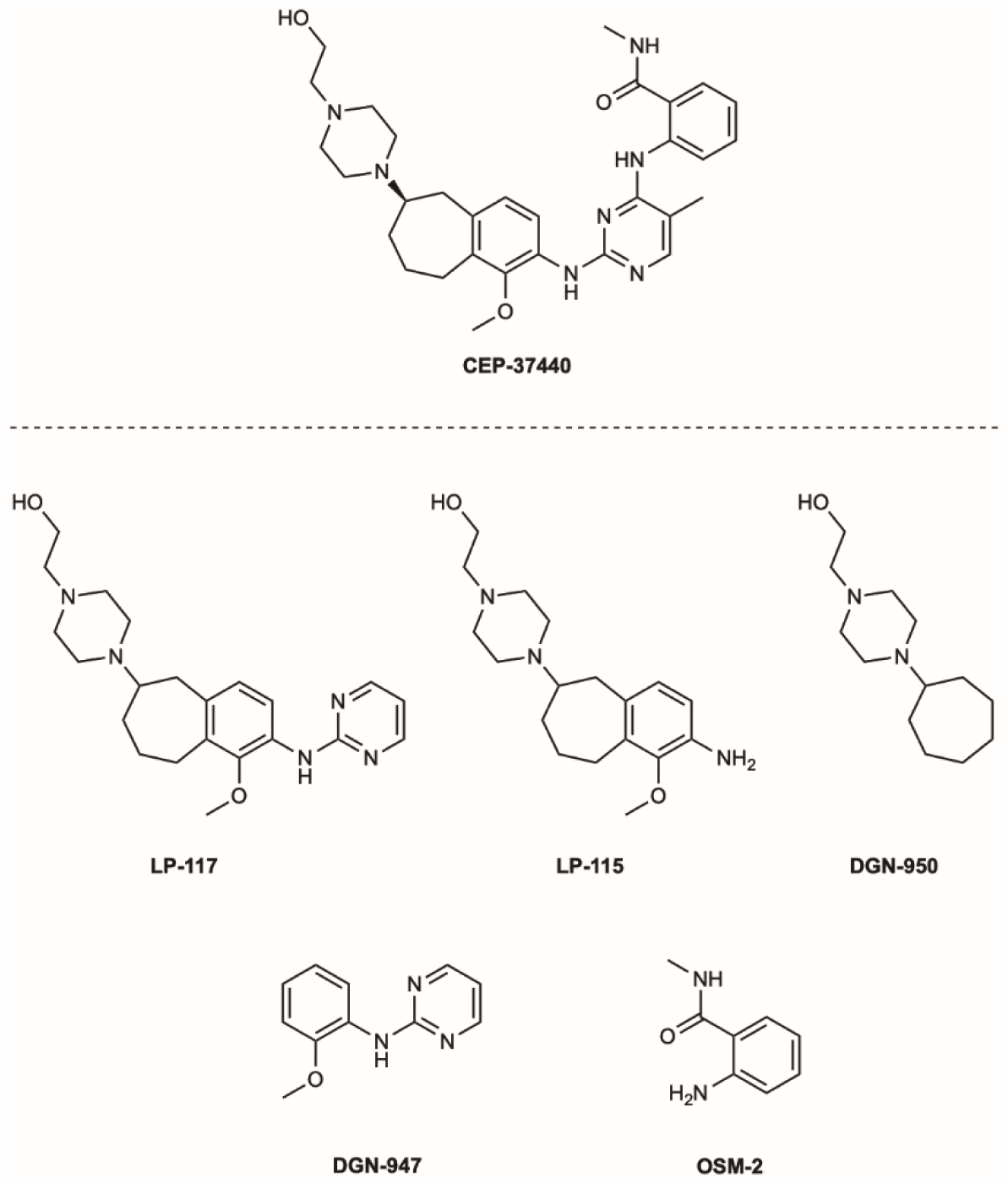
Deconstruction of the efflux pump-binding kinase inhibitor CEP-37440. Chemical structures of CEP-37440 and derived fragment-like substructures.

All fragments were assessed for TolC binding using MST. Compounds OSM-2, DGN-950, and DGN-947 did not interact with the OMF up to a concentration of 16 mM, whereas LP-117 appeared to weakly interact with the protein (Fig. S3). Compound LP-115, a substructure of LP-117, bound to TolC with a low millimolar affinity (K_D_ = 3.5 mM [95% CI: 1.7; 7.8]; Fig. 3a). Additional measurements showed no significant interaction with AcrA, and a weak binding to AcrB (K_D_ = 1.5 mM [95% CI: 0.80; 2.9]; Fig. 3a). However, LP-115 did not show antimicrobial potentiating activity in bacteria tested up to a concentration of 125 µM (Table S2).

**Fig. 3.**
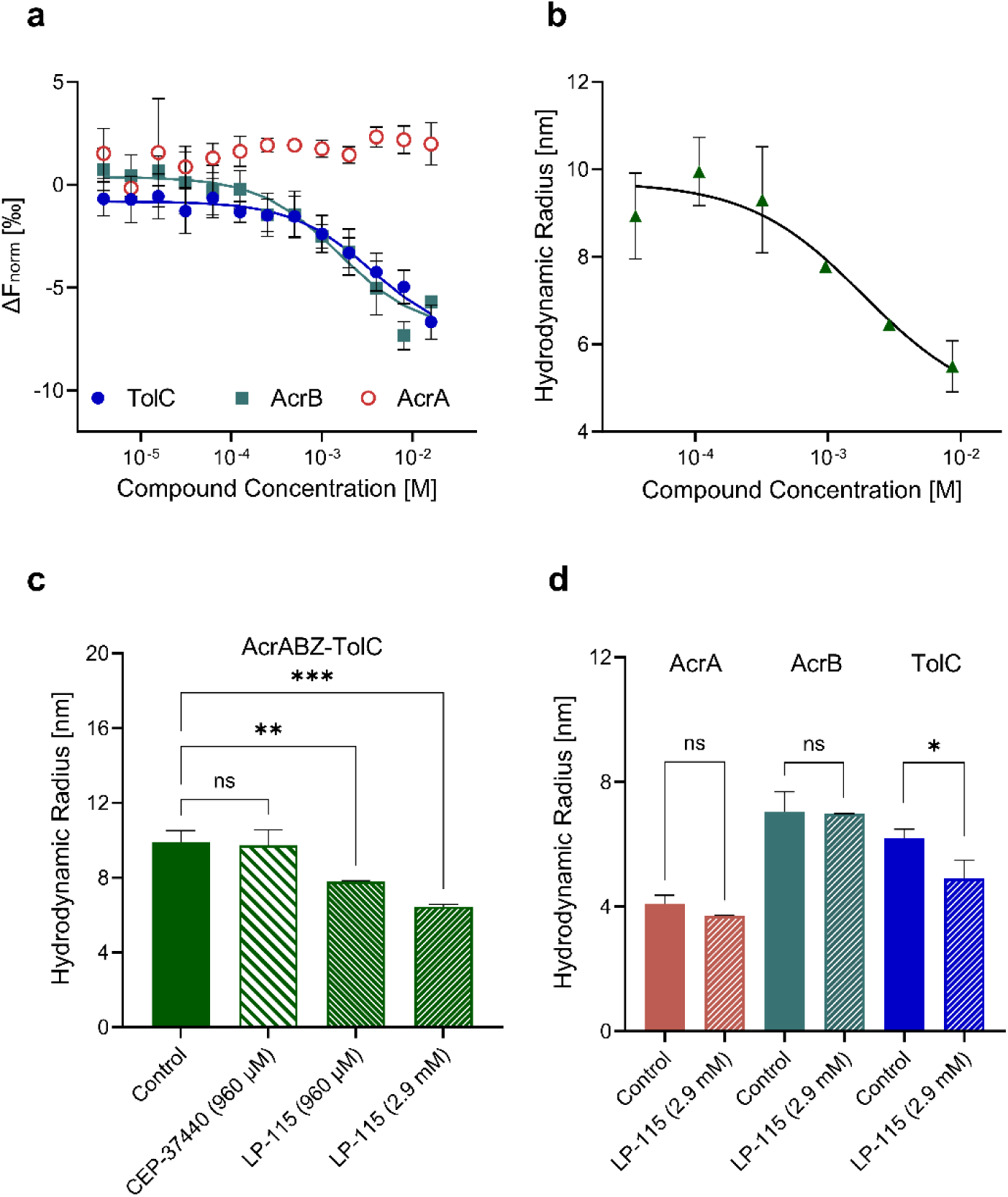
Compound-mediated disruption of the AcrABZ-TolC efflux pump. **a** LP-115 binding studies with AcrA (red), AcrB (petrol) and TolC (blue), using microscale thermophoresis at 0-1 s laser-on-time. Data points are presented as mean and SD and are derived from Fig. S5. **b** Reduction of the hydrodynamic radius of the AcrABZ-TolC efflux pump at various LP-115 concentrations as determined by dynamic light scattering. Hydrodynamic radii are derived from the raw data in Fig. S6 using Raynals webtool. **c** Hydrodynamic radius of the AcrABZ-TolC complex at 960 µM concentrations of CEP- 37440 and LP-115 as well as at 2880 µM of LP-115. Hydrodynamic radii are derived from the raw data in Fig. S7 using Raynals webtool. **d** Hydrodynamic radius of isolated efflux pump proteins in absence and presence of 2880 µM LP-115. Hydrodynamic radii are derived from the raw data in Figs. S8 and S9 using Raynals webtool. Data is presented as mean ±SD (n=3). Differences in hydrodynamic radius of protein alone and in presence of either CEP-37440 or LP-115 were analyzed by two-sided t-test. *p < 0.05; **p < 0.01; ***p < 0.001.

To further investigate the interaction of LP-115 and CEP-37440 with the entire efflux pump AcrABZ-TolC, dynamic light scattering (DLS) measurements were conducted. The complex was covalently linked between the AcrB and AcrA subunits, whereas TolC and AcrZ interacted non-covalently with the other subunits. LP-115 decreased the hydrodynamic radius of AcrABZ-TolC (9.9 nm ±0.5 nm) in a concentration-dependent manner (Figs. 3b and 3c). In contrast, CEP-37440 did not alter the hydrodynamic radius of AcrABZ-TolC at 960 µM (Fig. 3c). DLS measurements of LP-115 with AcrA or AcrB did not indicate any change in the hydrodynamic radius of the proteins at the corresponding concentrations; however, the substance decreased the hydrodynamic radius of TolC (Fig. 3d).

### Cryo-EM studies of LP-115 bound to AcrABZ-TolC

To confirm that TolC is the target of LP-115, a high-resolution structure of the AcrABZ-TolC efflux pump complexed with LP-115 was determined using single-particle cryo-EM (Figs. 4, S10–S12; Table S3). The complex was prepared detergent-free using the NSPr peptidisc method^21^, providing a more native-like environment for structural analysis (Fig. S10). The global resolution of the cryo-EM map reached 3.31 Å, allowing for clear definition of the protein components (Figs. 4, S11c, and S12a). Although we identified density potentially attributable to LP-115 at the periplasmic tip of TolC, the local resolution was relatively low, likely due to inherent dynamicity in this region. To enhance the resolution of the inhibitor density, signals from AcrB and most of AcrA, which were not directly involved in the interaction with TolC, were subtracted from the particles. This was followed by local classification to decrease structural heterogeneity and improve alignment (Figs. 5, S11, and S12b). A locally refined map at 4.72 Å subsequently allowed for a more accurate fitting of the inhibitor density (Figs. 5a, 5b, S11e, and S12b). However, some ambiguity remained in the compound density, potentially resulting from the resolution limits in the C1 map, partial occupancy, and/or variability in ligand conformations. Consequently, the best fitted model of the AcrA-TolC-LP-115 complex was analyzed to elucidate the precise molecular interactions at the binding site and to gain insights into the structural and functional dynamics of the complex.

**Fig. 4.**
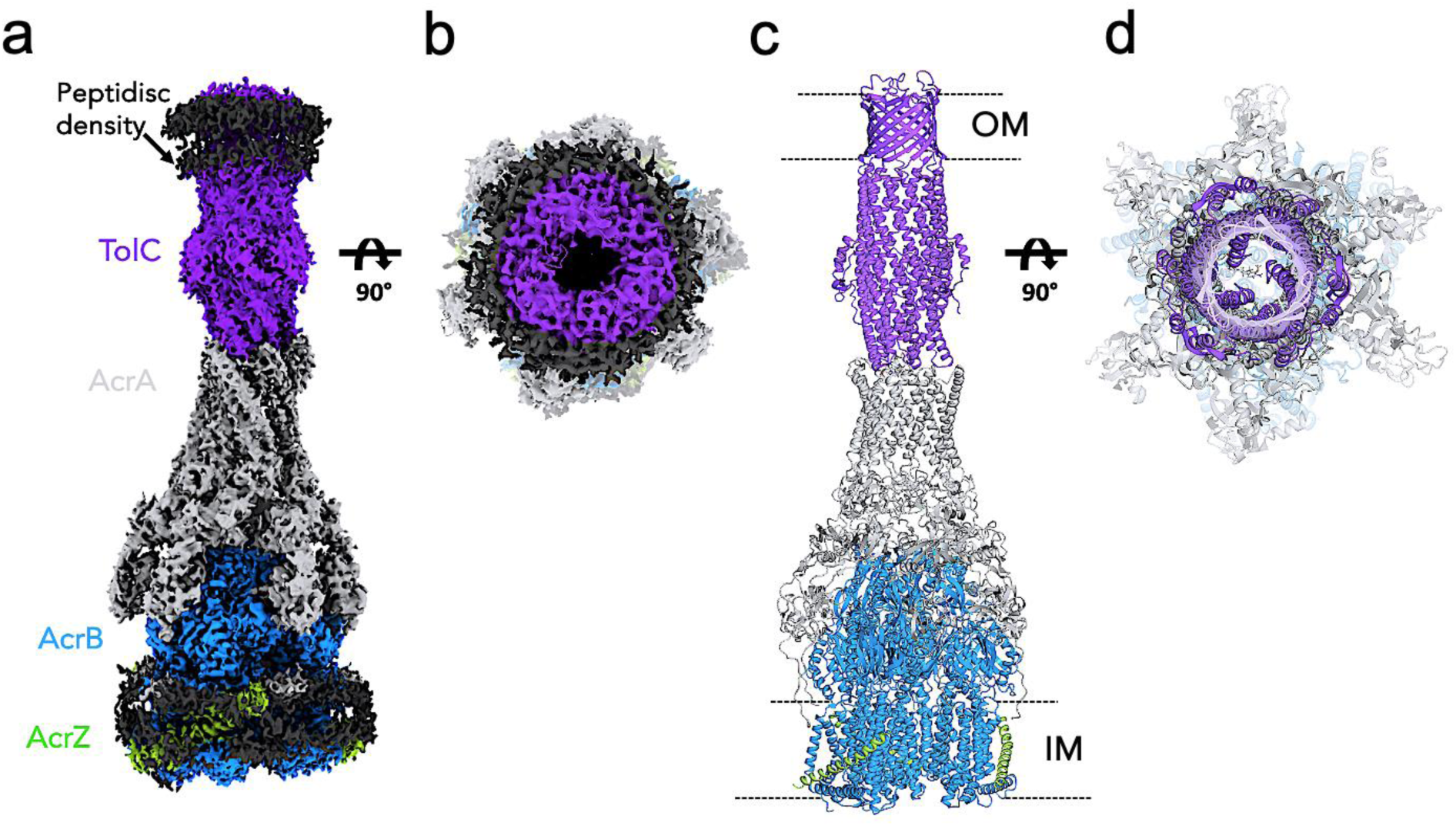
LP-115 blocks AcrABZ-TolC by binding to the periplasmic tip of the TolC-AcrA interface. **a-b** Cryo-EM structure of the efflux pump viewed through the membrane plane and from the outer membrane. **c-d** The atomic model of AcrABZ-TolC (in grey, blue, green and purple, respectively) bound to LP-115 (grey) is shown in ribbon representation as viewed through the membrane plane and from the outer membrane (PDB ID: XXXX).

**Fig. 5.**
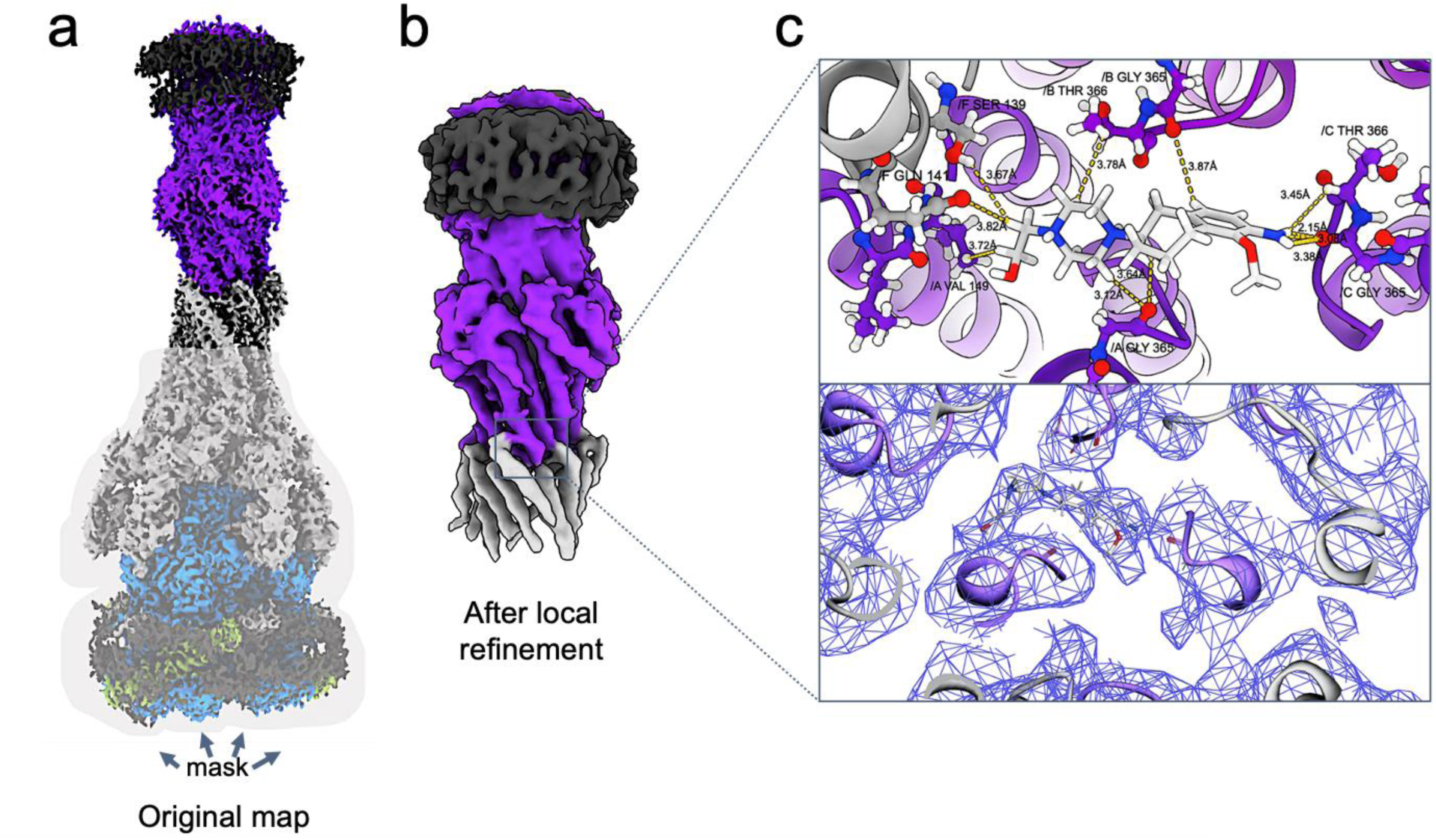
Cryo-EM analysis of LP-115 binding in the TolC-AcrA interface. **a** Cryo-EM structure of the efflux pump showing mask as transparent grey surface used for local refinement focusing on the LP-115 binding site. **b** The cryo-EM map after local refinement showing the TolC (purple) and AcrA (grey) subcomplex. **c** Upper panel: atomic model of LP-115 binding environment in the TolC-AcrA interface, showing amino acids (grey and purple sticks) forming hydrogen bonds and some weaker interactions (yellow dotted lines). Lower panel: electron density map of LP-115 binding environment.

The cryo-EM structure revealed LP-115 to bind at the interface of TolC (chains A–C) and AcrA (chains D–I) of the efflux pump complex (Fig. 5c). The amine group of LP-115 forms a hydrogen bond with Gly365 of chain C. In addition, several non-polar interactions are present involving Val149 (chain A), Gly365 on chains A and B as well as Thr366 on chain B. Furthermore, LP-115 interacts with with AcrA via non-polar interactions between the hydroxyethyl substituent of the LP-115 piperazine ring and the side chains of Ser139 (chain F) and Gln141, both located on chain F.

The structural details revealed by the binding of LP-115 to the AcrABZ-TolC complex show that the TolC-AcrA interface closely resembles a closed state of the system. The periplasmic tip of the TolC channel, where LP-115 binds, exhibits a dense clustering of residues within the homotrimer and notable constriction at the AcrA-TolC interface, as shown in Figs. 4 and 6. This arrangement is consistent with previously reported closed state cryo-EM structures^22^. In AcrA, which is typically seen in two distinct conformational states within the hexamer in its apo form, the binding of LP-115 disrupts the intermolecular interactions that maintain these conformations, leading to larger gaps between the AcrA protomers (Fig. 6a) and altering or disrupting the stabilizing bonds that maintain the two conformers. Furthermore, gaps are observed at the interfaces between adjacent AcrA dimer pairs, preventing the helical hairpin, lipoyl, and β-barrel domains of AcrA from tightly packing to seal the channel (Figs. 4, 6a, and 6b). These gaps must be closed during the transport process to prevent substrate leakage into the periplasm. These global conformational changes underscore the structural instability induced by LP-115 binding and its significant impact on the connectivity of the TolC-AcrA interface.

**Fig. 6.**
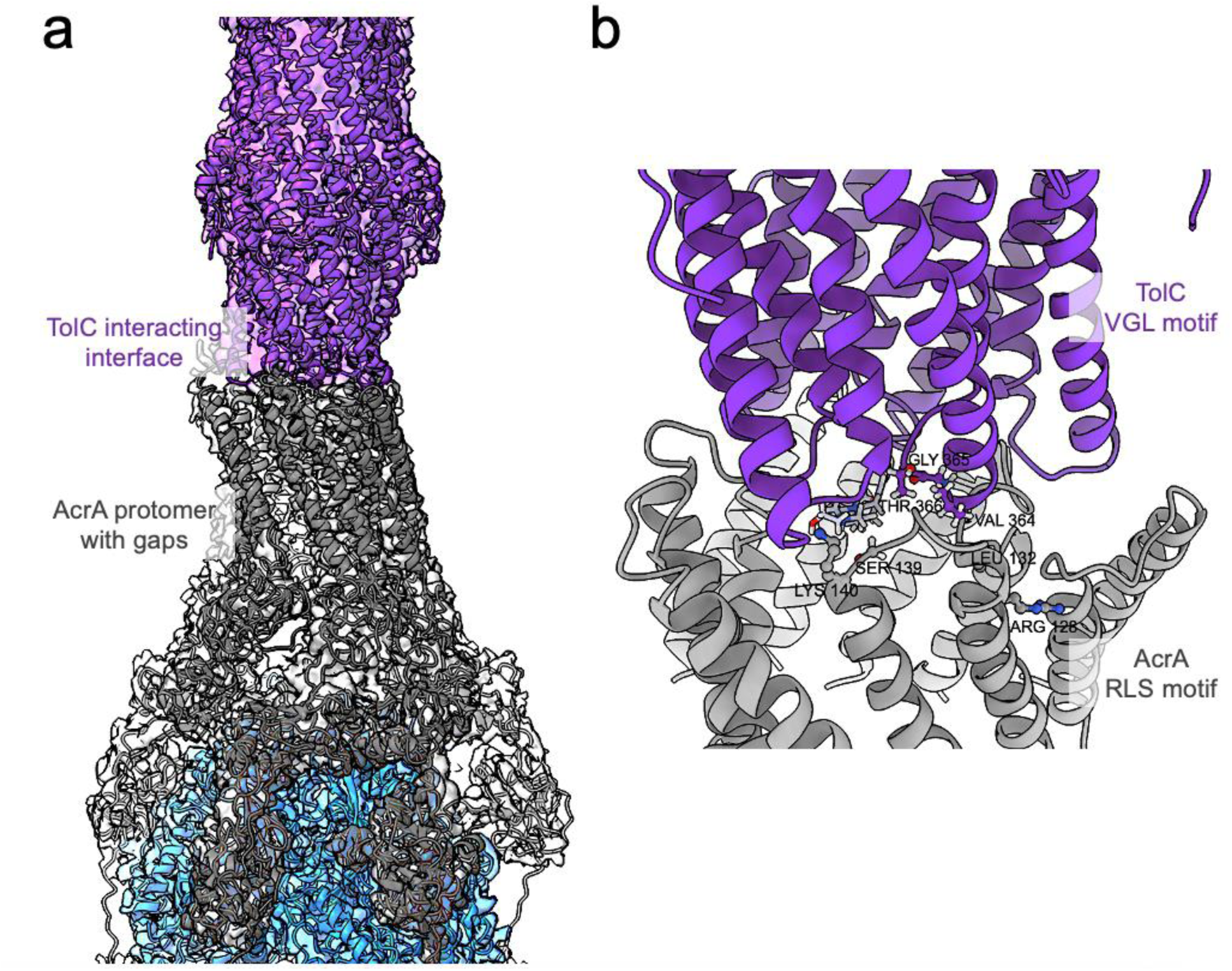
Disrupted interactions and conformational changes between TolC and AcrA in the AcrABZ-TolC pump bound to LP-115. **a** The cryo-EM density map at 3.1 Å resolution of the efflux pump reveals the detailed architecture at the interaction site. The fitted model highlights the tip-to-tip contacts between the TolC protomers (purple) and the α-helical hairpins of AcrA (grey), demonstrating their molecular assembly in the closed state. **b** A closer examination of the tip-to-tip interface between TolC and AcrA is presented. At this interface, residue pairs of the conversed VGL/T motif of TolC and RLS motif of AcrA are shown (purple and grey sticks).

The binding site of LP-115 is determined by the interaction between the conserved motifs of AcrA and TolC, which are pivotal for their association (Fig. 6b). The TolC protein features a conserved motif within the helix-turn-helix tips of its coiled-coil α-hairpin, known as the VGL/T motif (Val-Gly-Leu/Thr), which is essential for direct interaction with AcrA through AcrA’s conserved RLS motif (Arg-Leu-Ser) for the active transport process^23,24^. Specifically, at the interface between AcrA protomer-I and TolC in the open state, the interaction involves the backbone of Gly365 from the VGL/T motif of TolC and the backbone of Lys140 alongside the side chain of Ser139 from AcrA’s RLS motif^7,23^. Moreover, in the LP-115-bound complex, the contacts between Thr366 of TolC (VGL/T motif) and Leu132 of AcrA (RLS motif), which are critical for channel opening, are disrupted^7,23^.

## Discussion

Efflux pump inhibition is a promising strategy for improving antibiotic efficacy. Several options for inhibiting drug efflux from Gram-negative bacteria exist, for example suppression of efflux pump expression or inhibition of the efflux pump function, either by ligand-mediated blockage or disruption of efflux pump complex formation^2,25^. So far, most efflux pump inhibitors target AcrB, for example by binding to the deep pocket such as MBX2319 or an allosteric site in its transmembrane domain (BDM88855)^8,26^.

TolC, and its homologs in other Gram-negative bacteria, have been out of scope as potential drug targets for many years and all efforts to develop efflux pump inhibitors concentrated on AcrA and AcrB^27^. Very recently, a study on a peptide targeting TolC has shown a beneficial effect in combination with antibiotics, thereby suggesting the suitability of TolC as drug target^12^. Addressing outer membrane factors, such as TolC in *E. coli* or OprM in *Pseudomonas aeruginosa,* with small molecules or peptide-based therapeutics offers several advantages over targeting other efflux pump components. Depending on the binding site, TolC-addressing compounds may not require crossing the outer membrane, which represents a major hurdle for many small molecule drugs targeting Gram-negative bacteria^28^. Furthermore, TolC is part of different efflux pump complexes (e.g. HlyBD, MacAB, EmrAB, EmrKY, AcrAB, AcrAD, OqxAB, and MdtEF) most of which are known to accept at least one antibiotic as substrate. Consequently, a drug specifically addressing TolC could be administered as a broad-spectrum adjuvant for antibiotics.

In this study we identified the kinase inhibitor CEP-37440 to act as efflux pump inhibitor. The compound reduces the extrusion of the efflux pump substrate Hoechst 33342 and potentiates the antibacterial activity of piperacillin and levofloxacin, while it does not exhibit any intrinsic antibacterial potential and synergistic activity on an *E. coli* mutant lacking a functional AcrAB-TolC pump. Binding studies with isolated efflux pump proteins revealed the compound to weakly interact with AcrB and TolC, thus supporting the hypothesis that the synergistic effect of CEP-37440 in combination with antibiotics is mediated via interaction with the AcrAB-TolC pump. CEP-37440 has been described to selectively inhibit focal adhesion kinase and anaplastic lymphoma kinase^29,30^. Due to its moderate synergistic effect with piperacillin, the large molecular weight, and the weak TolC affinity and selectivity, the compound does not represent a promising starting structure for the development of a TolC-targeting antibiotic adjuvant.

Hit deconstruction appears to be an alternative approach to identify substructures of weakly potent compounds or substances with limited potential for further optimisation^31^. Resulting fragments with modest or low binding affinity or biological activity might be further developed using fragment-growing strategies. Thereby, hit deconstruction can be considered as link from classical drug discovery to fragment-based drug design.

The CEP-37440 fragment LP-115 showed very weak TolC affinity and selectivity as it bound to AcrB with a similar K_D_ compared to TolC. This is not surprising as AcrB is known to bind structurally diverse compounds^32^. In contrast to CEP-37440, LP-115 was capable of disassembling the AcrABZ-TolC complex as shown by the reduction of the hydrodynamic radius using DLS. The cryo-EM data enabled a more detailed view into the structural changes induced by the compound. The structural data suggest that the interaction of LP-115 with the TolC VGL/T motif and the AcrA RLS motif obstructs formation of key interactions necessary for transitioning to the open conformation and forces the TolC–AcrA interface into a closed-state configuration with notable gaps in between the AcrA protomers. Considering the DLS results showing a ligand-mediated disassembly of the complex, it appears plausible that the observed conformational changes in the cryo-EM structure represent a transitional state from the assembled to the disassembled form.

Based on the LP-115 binding mode, possible compound optimisation can be suggested that presumably strengthen the interaction with interfacial amino acids and may improve the efficacy of the substance. This may involve modifications that enable hydrogen bond formation with the polar side chains of nearby located amino acids such as Glu365 (TolC, chains A & B) (AcrA), Ser139, or Gln141 (AcrA).

The low binding affinity as well as the ambiguity in compound density and the limited number of protein-ligand interactions seen in the cryo-EM structure suggest a highly dynamic interaction of the compound with the TolC–AcrA interface. Due to the absence of a defined binding pocket at this location, the compound is largely solvent-exposed, including hydrophobic areas which is energetically unfavorable. Binding of an asymmetrical compound to a symmetrical site is not optimal, which might serve as a further guideline for compound optimisation efforts.

Despite most efflux pump inhibitor development efforts concentrate on classical AcrB inhibitors, several efflux pump assembly inhibitors have been reported in the last years. A designed ankyrin repeat protein (DARPin) has been shown bind to AcrB, thereby inhibiting the interaction with the periplasmic adapter protein AcrA^33,34^. Moreover, NSC60339 has been demonstrated to interfere with AcrA and its assembly in the pump, presumably by inhibiting its hexamer formation^10^. More recently, the compound has been proven to rigidify AcrA and is expected to interfere with the communicating role of AcrA between AcrB and TolC^35^.

Our data allow for the assumption that the TolC–AcrA interface is amenable to disruptions by small molecules capable of addressing the VGL/T and RLS motif of both proteins. Therefore, disruption of the AcrA–TolC interaction represents a promising mechanism of action for efflux pump inhibitors. LP-115 provides the basis for the development of potent compounds targeting this previously unexplored protein-protein interaction site. It remains to be elucidated whether compounds interfering with the AcrA–TolC interface are capable of disrupting other TolC-containing efflux pumps in *E. coli*, for example MacAB-TolC and thus offer broad efflux pump inhibition.

## Methods

### Molecular Modeling

Three-dimensional coordinates and dominant tautomers for an internal library of approximately 6500 approved and investigational drugs were generated using the molecular modeling suite MOE (Molecular Operating Environment) version 2019.01.02 (Chemical Computing Group Inc., Montreal, Canada). The library was filtered for compounds containing a positively charged nitrogen atom, resulting in a subset of 1216 substances. X-ray crystal structures for *E. coli* TolC (PDB IDs 1EK9 and 1TQQ) were downloaded from the Protein Data Bank^6,13,36^. All non-protein atoms were removed. Both structures were superposed on Cα atoms using MOE. Protein structures were prepared using Protonate3D in MOE and energy minimized (Amber14:EHT force field with R-Field implicit solvation model) using a three-step procedure (Cα atoms, backbone, all atoms), each step including a tether of 0.5 and an RMS gradient of 5. Compounds were docked into the periplasmic tip area of both structures using GOLD (Cambridge Crystallographic Data Centre, Cambridge, UK). The search space was defined by a sphere of 15 Å radius centered within the small opening at the periplasmic tip. For each compound 100 docking runs were conducted. The early termination option was switched off.

### Bacterial strains and reagents

The clinical control strain *Escherichia coli* BW25113 (parental strain) and the single-gene knock-out mutant strain *E. coli* JW5503 (ΔtolC) were obtained from the *E. coli* collection of the National BioResource Project at the National Institute of Genetics (Japan)^37^. Piperacillin sodium salt, phenylalanine-arginine β-naphthylamide dihydrochloride (PAβN) and Hoechst 33342 trihydrochloride (H33342) were purchased from Sigma. Selected compounds were purchased from Biomol (CEP-37440, methapyrilene), SCBT (ICI- 89406), Selleckchem (diperodon), Sigma-(BW723C86, mirabegron, zolmitriptan), and Tocris (xamoterol).

### H33342-based accumulation assay

The assay was performed as previously described^38^. Compounds were added to the bacterial suspension to achieve a final concentration of 100 µM. As positive control PAβN was used at 50 µM. *E. coli* JW5503 cells can naturally and rapidly accumulate H33342 intracellularly, thus showing a maximum fluorescence. The accumulation obtained in *E. coli* BW25113 treated with compounds and PAβN was compared to the accumulation observed in JW5503 and calculated as percentage values (*E. coli* JW5503 was assumed as 100 %). Intracellular H33342 was monitored using a multimode microplate reader (VarioskanLUX, Thermo Fisher Scientific) to measure the fluorescence (ex 355 nm, em 460 nm) of the wells at 37 °C every 5 minutes for 30 minutes. The experiment was performed once with two technical replicates for each compound. Further details are available in Supplementary Information.

### MIC-based assay for potentiation of antibiotics and antibacterial activity of compounds

MIC assays were performed by the broth microdilution method in 96-well plate format according to the Clinical and Laboratory Standards Institute (CLSI) guidelines^39^. For the potentiation activity, an equal volume of bacterial suspension and compound solution, diluted into Cation Adjusted Mueller Hinton Broth (CAMHB) containing piperacillin (final concentration of 1 µg/mL) or levofloxacin (final concentration of 0.01 µg/mL) were mixed in a 96-well plate and plates incubated at 37 °C for 24 hours. The intrinsic MIC of each compound against *E. coli* BW25113 was determined using a similar assay, with the exception that the serial dilutions of compounds in CAMHB did not contain antibiotics. PAβN at 100 µM was used as positive control in potentiation assays, while piperacillin or levofloxacin alone (4 µg/mL and 0.04 µg/mL, respectively) in MIC determination of compounds. Compounds were tested at concentrations ranging from 125 to 0.39 µM. Two independent experiments with three technical replicates each were carried out.

### Construction of vectors for TolC-3xFLAG-AcrZ-6xHis and disulphide-linked AcrA-AcrB for overexpression

The gene encoding the TolC protein was engineered with 3xFLAG epitopes prepared by two successive PCR reactions. First, two fragments from the wild-type TolC gene were amplified and used as templates for the insertion. For the first PCR reaction, the primers pEPF1w-TolC-for and pEPReFt-3xFLAG-rev were used and for the second PCR reaction the primers pEPF1w-TolC-rev and pEPReFt-3xFLAG-for (Table S4). The PCR products were resolved on 1% low melting point agarose gel (Sigma) and the bands were excised. The excised bands were then heated at 70 °C to melt the gel and mixed in one PCR reaction for amplification of the entire TolC gene consisting of the 3xFLAG insertion using primers pEPF1w-TolC-for and pEPF1w-TolC-rev (Table S4). The full-length TolC- 3xFLAG PCR amplified gene product was treated with restriction enzymes AvrII and NdeI (NEB) and resolved on 1 % low melting point agarose gel. The gel bands were excised and ligated with T4 ligase (NEB) into a pRSF-duet plasmid treated AvrII and NdeI (NEB) and dephosphorylated with calf intestinal alkaline phosphatase (CIP, NEB) following the manufacturer’s instructions. The engineered TolC-3xFLAG was digested with AvRII and NdeI enzymes from pRSF-duet plasmid and then subcloned into the second multiple cloning site (MCS) of expression vector PRSF-duet encoding for AcrZ-6xHis-tagged, resulting in the construct PRSF-duet-AcrZ-6xHis-TolC-3xFLAG. The AcrAB sub- assembly has been engineered to form a disulphide bond for stabilization during expression and purification procedures as the complex is highly prone to dissociation^22^. For this, the substitution S273C has been introduced in AcrA and S258C in AcrB by site-directed mutagenesis using pAcBH as template (Table S4)^22^.

### Protein expression of AcrABZ-TolC

Co-expression and cellular membrane preparation of the recombinant AcrABZ-TolC- 3xFLAG was prepared as previously described with some modifications^7^. The AcrABZ-TolC-3xFLAG components were expressed in *E. coli* strain C43 (ΔacrAB and ΔTolC). Cell pellets were resuspended in lysis buffer (150 mM NaCl and 50 mM Tris pH 7.5). After pelleting the membranes as previously described^7^, 4.5 g of cellular membrane extracts were solubilized with buffer A (400 mM NaCl, 50 mM Tris pH 7.5 and 1 % wt/vol n-dodecyl-β-D-maltopyranoside (β-DDM). The membrane pellet was solubilized for 3 hours at 4 °C by stirring and incubated with 0.5 ml of nickel-nitrilotriacetic acid (Ni-NTA) beads (Qiagen) for nickel affinity purification.

### Purification of AcrABZ-TolC-3xFLAG

Histidine tagged AcrABZ-6xHis-TolC-3xFLAG was purified by nickel affinity using Ni-NTA resins. The supernatant was removed, and the beads loaded onto a gravity flow-column. The column was washed with 10 column volumes (CVs) of buffer B (200 mM NaCl, 50 mM Tris pH 7.5 and 0.02 % wt/vol β-DDM). The AcrABZ-TolC-3xFLAG protein complex was eluted with buffer B + 0.5 M imidazole. Eluate fractions were verified by a 4–12% sodium dodecyl sulfate-polyacrylamide gel electrophoresis (SDS-PAGE) gel and then pooled together for a second purification step using anti-FLAG M2 affinity beads (Sigma). The pooled fractions were incubated with 0.5 mL anti-FLAG resin for 1 hour at 4 °C with gentle stirring. The column was washed with 5 CVs buffer B and 3 CVs buffer C (200 mM NaCl, 50 mM Tris pH 7.5) to gradually decrease the detergent critical micelle concentration (CMC) of the solubilized pump. Next, 2 mg/mL of assembled NSPr peptidisc^21^ in 20 mM Tris-HCl, pH 8.0 peptide solution was loaded onto the column and the beads were incubated for 30 minutes at room temperature with gentle shaking. This step was repeated one more time. Finally, the reconstituted pump free of detergent was eluted using buffer C plus 0.5 mg/mL of 3xFLAG peptide for the first elution step, and thereafter with buffer C alone. The fractions were verified with a 4–12% gradient SDS-PAGE gel and concentrated to 5–6 mg/mL using a Vivaspin column (MWCO: 100 kDa).

### AcrA, AcrB and TolC protein expression and purification

AcrA and AcrB were expressed and purified as previously described (Szal 2023)^41^. A pET24a-based plasmid for the expression of a C-terminally His-tagged full-length TolC was kindly provided by Mathias Winterhalter, Constructor University, Bremen, Germany.

*E. coli* C43(DE3)ΔAcrAB bacteria were transformed with the TolC expression plasmid (pET24a) were cultured in TB medium with 50 µg/mL kanamycin (Carl Roth GmbH + Co KG, Karlsruhe, Germany) at 37 °C and 200 rpm. At an OD_600_ of 1.3 the protein production was induced with isopropyl-β-thiogalactoside (IPTG; Biosynth AG, Staad, Switzerland) and shaken over night at 20 °C and 200 rpm. Pelleted cells were resuspended in 4 mL of 20 mM Tris/HCl pH 7.4, 150 mM NaCl with 0.25 mg/mL Lysozyme, 0.1 mg DNase I (AppliChem GmbH, Darmstadt, Germany) per 1 g cell pellet, and 1 tablet protease inhibitor cocktail (cOmplete™, Sigma Aldrich, Merck KGaA, Taufkirchen, Germany) per 100 mL resuspended cell pellet. Subsequently, the solution was lysed using a LM10 Microfluidizer (Microfluidics). After pelleting the cell debris (5000 x g, 4 °C, 15 min), membranes were pelleted by centrifugation at 137,000 x g for 1h at 4 °C. Membranes were resuspended and solubilized in loading buffer (20 mM Tris/HCl pH 7.4, 150 mM NaCl, 10 mM Imidazole/HCl pH 7.5) at a concentration of 10 mL/g pellet in the presence of 1% DDM (CN26, Carl Roth GmbH + Co KG, Karlsruhe, Germany). The solubilized membranes were cleared by centrifugation at 137,000 x g for 1h at 4 °C. TolC was purified from the supernatant using a two-step chromatography approach on an ÄKTA system starting with an affinity chromatography step using a 5 mL HisTrap HP column (Cytiva) and followed by size exclusion chromatography step using a S200 PG 16/600 (Fig. S13). All chromatography steps were performed in the presence of 0.03% DDM. The protein was checked for identity and purity conducting an SDS-PAGE, Western Blot analysis (Fig. S14). Structural integrity was confirmed by nDSF measurements using a NanoTemper Tycho NT.6. The effect of TolC protein storage at 4 °C between the measurements was investigated using comprehensive mass spectrometric analysis (Fig. S15).

Aliquots of TEVp were kindly provided by the Protein Production facility at CSSB Hamburg.

### Microscale thermophoresis (MST)

Label-free MST (n=5) with AcrA, AcrB, and TEVp was carried out using a Monolith NT.115 (NanoTemper Technologies) as previously described^40^. Measurements with TolC were conducted in phosphate-buffered saline (PBS, pH 7.4) with 0.03 % DDM. The concentration of TolC was set to 5 µM, while the concentration of AcrA and TEVp was set to 5 µM and of AcrB to 1.25 µM. CEP-37440 was titrated in a three-fold dilution series in the respective buffer. The final DMSO concentration was 4 % which did not denature TolC (Fig. S16). The solutions were incubated at room temperature for 15 minutes to reach equilibrium state and measured in Monolith LabelFree Premium Capillaries (NanoTemper Technologies).

The fragments were measured with labeled MST using the instrument’s red filter set. TolC, AcrA and AcrB were labeled with the Protein Labeling Kit RED-NHS 2nd Generation (NanoTemper Technologies) in their respective buffers. Labeling related structural changes could be excluded by performing comparative nDSF measurements using a Tycho NT.6 (NanoTemper Technologies, Fig. S17). The degree of labeling was analyzed spectrophotometrically using a NanoDrop® ND-1000 UV-Vis spectrophotometer (Thermo Scientific GmbH, Table S5). The parameters (temperature, on- and off-times, IR-laser power) and buffers were not changed for the labeled MST. The LED power was set to 20% and the fragments were diluted in a two-fold dilution series. The protein concentration was set to 20 nM for all proteins. The protein-ligand-mixture was incubated for at least 15 minutes and loaded in Monolith Premium Capillaries (NanoTemper Technologies).

Data analysis was conducted with the ThermoAffinity web tool (https://spc.embl-hamburg.de)^41^ using a one-site fitting at 1 to 2 seconds MST signal for all compounds. Data points that differed by more than 20 % in the initial fluorescence or showed irregular MST traces were removed from MST evaluation as recommended by the NanoTemper Technologies. Compounds with curves that could not yield a 95 % confidence interval or an estimation of the parameters (RF1; RF2; K_D_) were interpreted as not binding.

### Dynamic light scattering (DLS)

DLS measurements were conducted with a Nanotemper Panta using Prometheus High Sensitivity Capillaries (NanoTemper Technologies). LP-115 was diluted in a 1:3-dilution series with 50 mM Tris pH 8.0, 150 mM NaCl and 0.02% DDM. The final concentration of AcrABZ-TolC was set to 78 nM, while the final DMSO concentration was 4%. For single point measurements with CEP-37440, the complex concentration, buffer and DMSO concentration were the same. For single point measurements with the subunits, the respective buffers were used as stated above and the protein concentration was set to 4 µM. All solutions were centrifuged before mixing (compound solution at 20,000 x g for 5 minutes and the complex at 3,500 x g for 10 minutes). The measurement was conducted in triplicate at 25 °C with 10 acquisitions in 3 scans at a wavelength of 405 nm, and a scattering angle of 147°. The analysis was conducted with Raynals web tool^42^ at the eSPC platform (https://spc.embl-hamburg.de/app/raynals), the regularisation α was fixed and set to 0.05. For the analysis of the concentration dependent complex destabilisation, a three parametric sigmoidal fitting using GraphPad Prism 9.2.0 (GraphPad Software) was utilized. A two-sided unpaired t-test was conducted for the statistical analysis of the hydrodynamic radii of untreated and treated (2880 µM LP-115) TolC, AcrA and AcrB, while an ordinary One-way ANOVA was conducted for the statistical comparison of untreated and treated (960 µM CEP-37440, 960 µM and 2880 µM LP115) AcrABZ-TolC.

The MST and DLS graphs were plotted using GraphPad Prism 9.2.0 (GraphPad Software).

### Cryo-EM sample preparation

For the AcrABZ-TolC pump-LP115 samples, 3.5 μL aliquots of purified protein in peptidisc (protein concentration = 0.9 mg/mL) were added to glow-discharged holey carbon grids (Quantifoil Au R1.2/1.3, 300 mesh). Excess sample was blotted away with an FEI Vitrobot (IV) (100% humidity, 4°C, blotting force ranging from −2 to +2, 3 s blot time). After blotting, the grids were vitrified in liquid ethane and screened on a FEI Talos Arctica. All datasets were collected on a FEI Titan Krios equipped with a Gatan K2 camera and the data collection parameters for the specimens are summarized in Table S3.

### Image processing

All processing steps were performed in cryoSPARC 4.1.2^43^. A schematic overview of the processing pipeline is displayed in Figure S11 and data collection and processing parameters are summarized in Table S3.

Several rounds of 2D classification were used to remove aberrant particles resulting to an initial 300,383 particle class that was subjected in three state classification using *ab initio* reconstruction. A final subset of 91,632 particles was subjected to multiple rounds of 3D homogeneous and non-uniform refinement generating an AcrAB-TolC-LP-115 bound reconstruction at a global resolution of 3.31 Å without application of symmetry restraints (C1). The resulting map quality for the full pump was high, but the quality of the inhibitor binding site was low. Consequently, a focused mask on TolC-AcrA interfaces was created and used for the local refinement of the corresponding region.

Local refinement with the focused mask resulted in a 4.72 Å reconstruction (C1) with improved map quality for the inhibitor binding site (Figs. S11 and S12). The final resolution of the maps was estimated by 0.143 cut-off of the Fourier shell correlation (FSC). Local resolution variations were estimated with the ResMap wrapper in Cryosparc using the two independent half-maps (Figs. S11 and S12)^44^.

### Model docking and refinement

To build the atomic models, a cryo-EM structure of the AcrABZ-TolC in the closed state (data not published yet) was fitted into the map density in UCSF Chimera v.1.15.0^45^.

Missing amino acids were built using Coot. In addition, Ramachandran outliers and overall geometry were manually fixed in Coot^46^. Models were refined with real-space refinement via Phenix v.1.20.1^47^ and ISOLDE v.16.0^48^. The quality of the stereochemistry was assessed using Molprobity^49^. For structure visualization and image creation ChimeraX v.1.6.1^50^ was used. The statistics for model refinement are summarized in Table S3.

## Data Availability

Atomic coordinates and maps reported in this paper have been deposited in the Protein Data Bank under accession number XXXX. Source data are provided with this paper.

## Acknowledgements

We acknowledge technical support by the SPC facility at EMBL Hamburg and by the PP facility, University Medical Center Hamburg-Eppendorf, Hamburg, Germany at CSSB Hamburg. We thank the Core Facility Mass Spectrometric Proteomics at the Medical Center Hamburg-Eppendorf (UKE), Maria Riedner and Sönke Harder (Technology Platform Mass Spectrometry, Universität Hamburg & UKE) for mass spectrometry-based protein identification and protein quality control (the MALDI rapifleX was supported by the DFG project 426788273, and the maXis ETD by the DFG project INST 152/859-1). This project was supported under the framework of the JPIAMR – Joint Programming Initiative on Antimicrobial Resistance. B.W. was supported by the German Federal Ministry for Education and Research (01KI1827A). A.J. was supported by the state education development agency (1.1.1.5/ERANET/19/03), C.D.C and P.T. were supported by the Academy of Finland (Grant no. 326261).

## Author contributions

P.T., A.J., J.V., B.F.L, and B.W. designed experiments. B.W. conducted virtual screening. T.S. carried out biophysical studies. C.D.C. designed and conducted microbiological assays. M.M., L.P., J.V, and A.J. performed the chemical synthesis. T.S. and F.Z.R. expressed and purified protein supported by P.L. and S.W. E.P. and B.F.L. determined cryogenic electron microscopy structures. T.S., E.P., and B.W. wrote the manuscript. All authors contributed to data analysis as well as manuscript review and editing. B.W., A.J., and P.T. obtained funding for this work.

## Additional Information

Correspondence and requests for materials should be addressed to Björn Windshügel.

