## Supplementary Information for "Disassembly of the *Escherichia coli* AcrABZ-TolC efflux pump by ligand-mediated disruption of TolC-AcrA interfacial contacts"

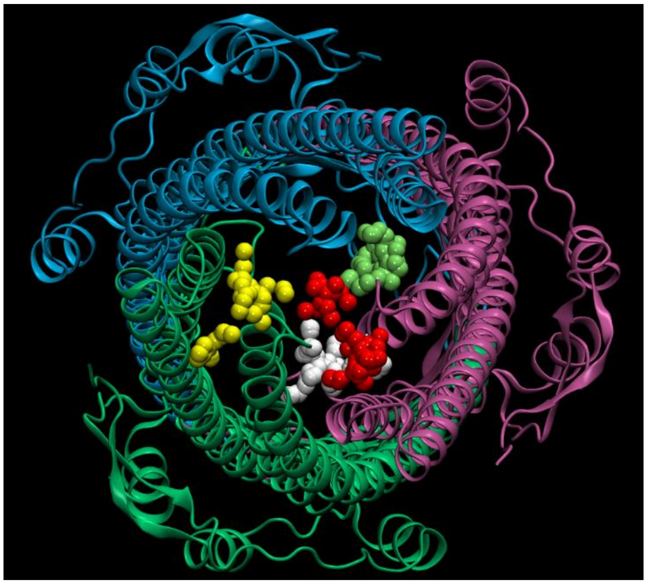

**Fig. S1 | Protein-ligand interaction possibilities at the periplasmic tip of TolC.** Potential binding sites for small molecules at the periplasmic tip of TolC as predicted by SiteFinder. Alpha spheres for different sites are shown in individual colors.

**Table S1 |** Compounds predicted to bind to TolC and selected for experimental validation.

| **Compound Name** | **Vendor** | **Catalogue #** |
|---|---|---|
| BW723C86 | Sigma Aldrich | B175-10MG |
| CEP-37440 hemifumarate | Biomol | Cay21278-5 |
| Diperodon HCl | Selleckchem | S4397 |
| ICI69406 | SCBT | sc-361212 |
| Methapyrilene 2 HCl | Biomol | Cay33434-100 |
| Mirabegron HCl | Sigma Aldrich | SML2480-5MG |
| Xamoterol | Tocris | 950 |
| Zolmitriptan | Sigma Aldrich | SML0248-10MG |

#### H33342-based accumulation assay

Bacteria were grown in Luria Broth (LB, HispanLab) for *E. coli* BW25113 and LB supplemented with 25 µg/mL of Kanamycin (Sigma) for *E. coli* JW5503, at 37°C for 24 hours under 200 rpm agitation. On the following day, overnight culture was inoculated into 5 mL of fresh LB and tubes incubated at 37° C at 200 rpm for about 2 hours until the logarithmic phase was achieved (i.e. McFarland scale of 2.5 - 3.0). Cells were harvested by centrifugation (4,000 x g, 5 minutes) and resuspended in 1 × phosphate buffer saline [137 mM NaCl (Riedel), 2.7 mM KCl (Riedel), 10 mM $Na_2HPO_4$ (J.T.Baker); and 2 mM $KH_2PO_4$ (Merck); pH 7.4, PBS]. Resuspended cells were then diluted (1:4) in PBS. Compounds were added to the bacterial suspension in order to achieve a final concentration of 100 µM. As positive control PAβN was used at 50 µM. Negative control constituted of PBS. Then, 180 µL of this mixture was added into 96-well black, clear bottom, microtiter plates (Perkin Elmer, Turku, Finland). Plates were incubated at 37 °C for 15 min, and then H33342 was added to each well (2.5 mM final concentration). Plates were then covered in foil, mixed for 5 minutes at 500 rpm, followed by kinetics fluorescence measurements.

**MIC-based assay for potentiation of antibiotics and antibacterial activity of compounds**

Bacterial colonies were taken from the Mueller Hinton agar (MHA, Neogen) overnight culture, inoculated into 0.9% saline solution and vortexed to ensure that the bacterial suspension was homogeneous. Bacterial suspensions were analysed using a densitometer (DEN-1, BioSan, Warren MI, USA) and adjusted to $1 \times 10^6$ colony forming units (CFU/mL) by diluting with cation adjusted Mueller Hinton broth (CAMHB, BD). An equal volume of bacterial suspension and test compound solution, diluted into CAMHB containing piperacillin or levofloxacin (final concentrations of 1 µg/mL and 0.01 µg/mL, respectively), were mixed together into plate wells (Thermo Fisher Scientific) and incubated for 24 hours at 37°C. Absorbance values measured at 600 nm using MultiskanGO plate reader (Thermo Fisher Scientific) were used for evaluating the antibacterial effects, by comparing to untreated controls, and expressed as percentage of growth inhibition. Minimum inhibitory concentration (MIC) values were defined as the lowest compound's concentration at which bacterial growth was inhibited by ≥90% compared to antibiotic- and compound-free control.

**Table S2 |** Inhibition of *E. coli* BW25113 and JW5503 growth by LP-115 in absence and presence of piperacillin.

| Compounds | ***E. coli* BW25113** | | ***E. coli* JW5503** | |
|---|---|---|---|---|
| | **MIC**[a] | **MPC4**[b] | **MIC** | **MPC4** |
| LP-115 | >125 µM (5.06 ± 12.79) | >125 µM (-25.18 ± 23.70) | >125 µM (0.19 ±13.85) | >125 µM (3.86 ± 8.65) |

[a]MIC: minimum inhibitory concentration was defined as the concentration of compound that inhibited *E. coli* growth by ≥90 %. If MIC was not achieved, the value is represented by the highest concentration of compound tested.

[b]$MPC_4$: minimum potentiating concentration was defined as the concentration of compound that decreased the MIC of antibiotic by 4-fold (i.e. piperacillin from 4 µg/mL to 1 µg/mL and from 0.25 µg/mL to 0.0625 µg/mL in *E. coli* BW25113 and JW5503, respectively). Experiments were performed once with three replicates each.

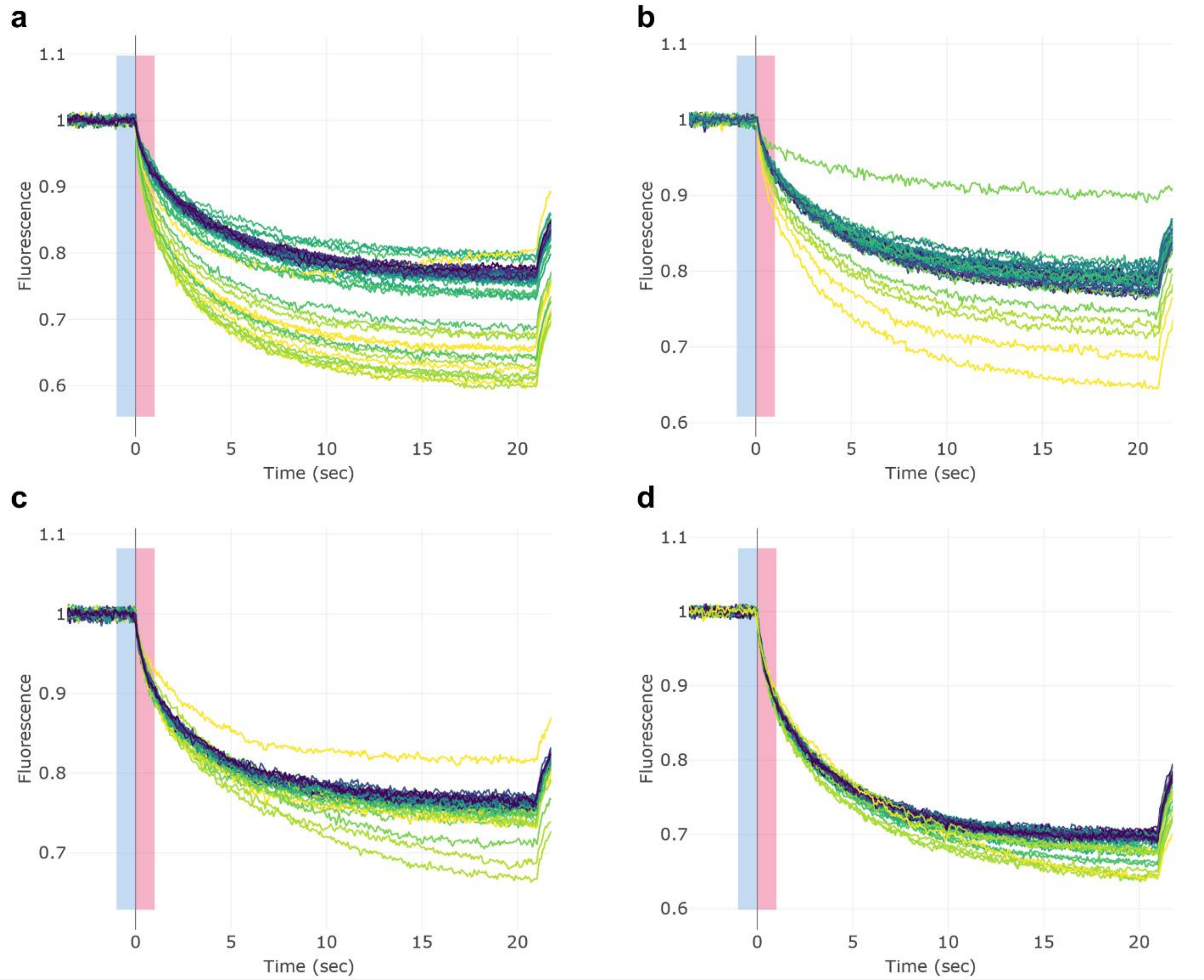


**Fig. S2 | Normalized MST Traces of CEP-37440 with the measured proteins. a** MST Traces of quintuplicate measurements of CEP-37440 with 5 µM TolC. **b** MST Traces of quintuplicate measurements of CEP-37440 with 1.25 µM AcrB. **c** MST Traces of quintuplicate measurements of CEP-37440 with 5 µM AcrA. **d** MST Traces of quintuplicate measurements of CEP-37440 with 5 µM TEVp. **a, b, c, d** The blue rectangle highlights the cold region, while the red rectangle highlights the chosen hot region. Traces with the highest compound concentration are highlighted in yellow (5.3 mM for TolC and TEVp; 1.7 mM for AcrA; 0.59 mM for AcrB), the second highest compound concentration in light green, the third highest concentration in green, followed by the lower concentrations in 7 to 10 shades of blue and with the lowest compound concentration in black (3.3 nM for TolC; 10 nM for AcrB, AcrA and TEVp).

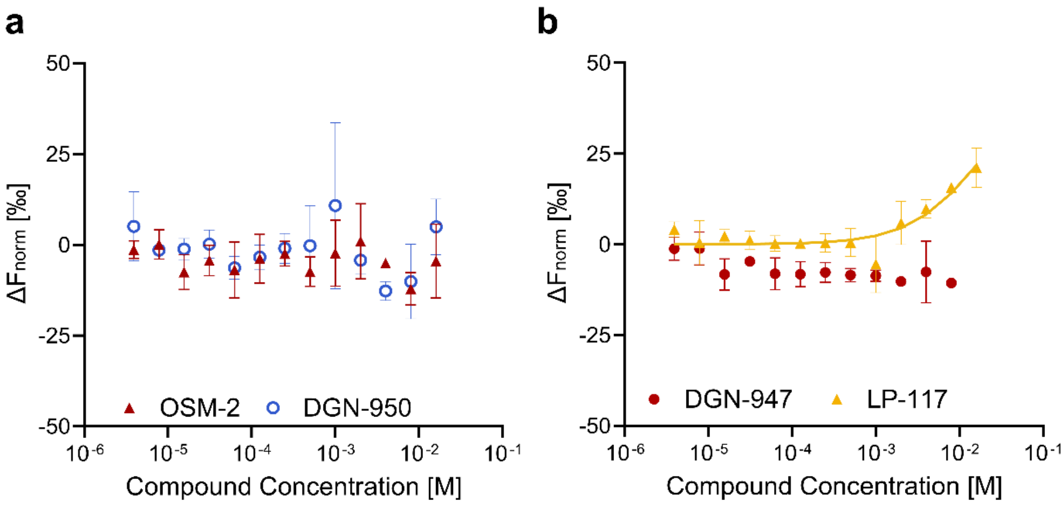


**Fig. S3 | Fragments interaction with different efflux pump proteins.** Binding studies of the CEP-37440 fragments **a** OSM-2 and DGN-950 **b** DGN-947 and LP-117 with fluorescently labeled TolC, using microscale thermophoresis at 19-20 sec laser-on-time. Data points are presented as mean ± SD and derived from Fig. S4.

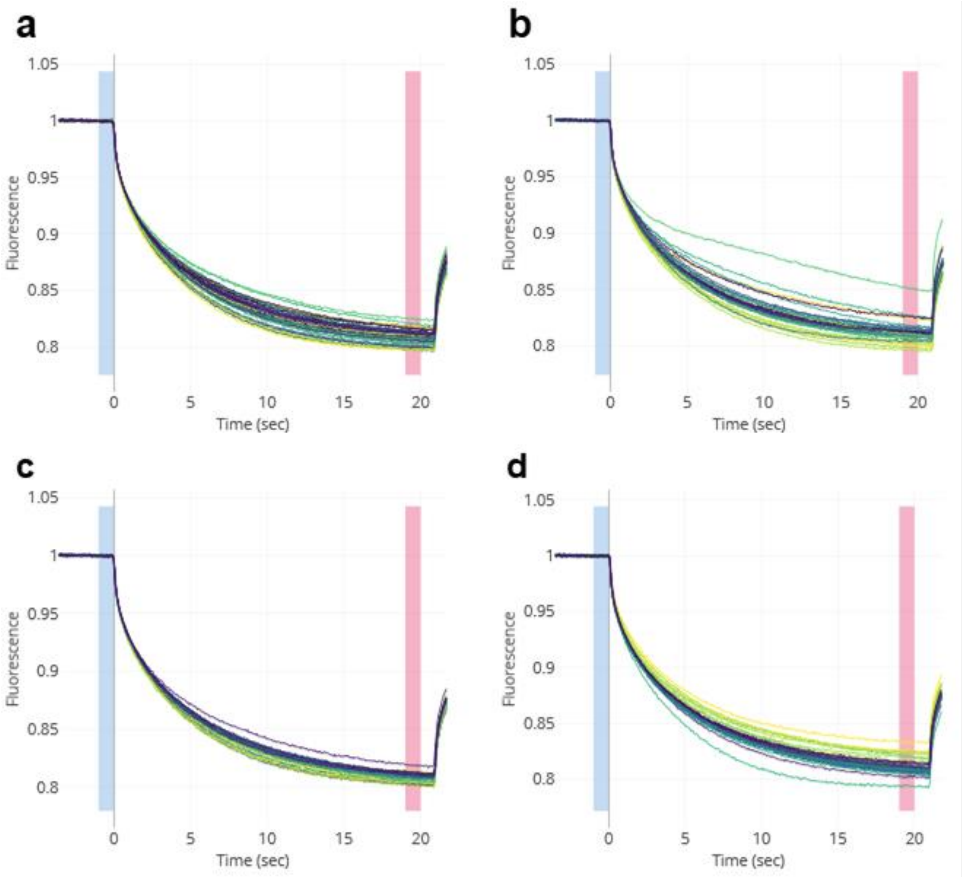


**Fig. S4 | Normalized MST Traces of the non-binding CEP-37440 fragments. a** MST Traces of triplicate measurements of OSM-2 with 20 nM TolC. **b** MST Traces of triplicate measurements of DGN-950 with 20 nM TolC. **c** MST Traces of triplicate measurements of DGN-947 with 20 nM TolC. **d** MST Traces of triplicate measurements of LP-117 with 20 nM TolC. **a, b, c, d** The blue rectangle highlights the cold region, while the red rectangle highlights the chosen hot region. Traces with the highest compound concentration are highlighted in yellow (16 mM for OSM-2, DGN-950, LP-117; 8 mM for DGN-947), the second highest compound concentration in light green, the third highest concentration in green, followed by the lower concentrations in 8 to 9 shades of blue and with the lowest compound concentration in black (3.9 µM).

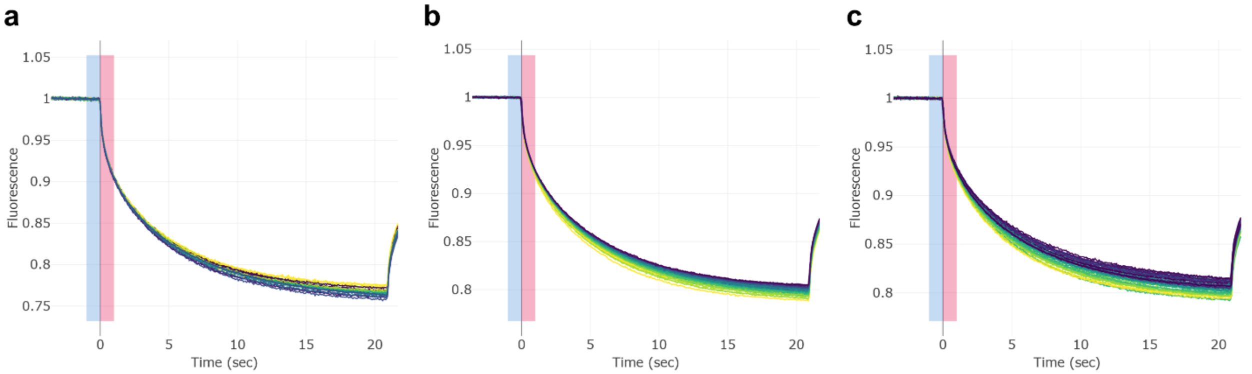


**Fig. S5 | Normalized MST Traces of LP-115 and the pump subunits. a** MST Traces of triplicate measurements of LP-115 with 20 nM AcrA. **b** MST Traces of quintuplicate measurements of LP-115 with 20 nM AcrB. **c** MST Traces of quintuplicate measurements of LP-115 with 20 nM TolC. **a, b, c** The blue rectangle highlights the cold region, while the red rectangle highlights the chosen hot region. Traces with the highest compound concentration are highlighted in yellow (16 mM), the second highest compound concentration in light green, the third highest concentration in green, followed by the lower concentrations in 9 shades of blue and with the lowest compound concentration in black (3.9 µM).

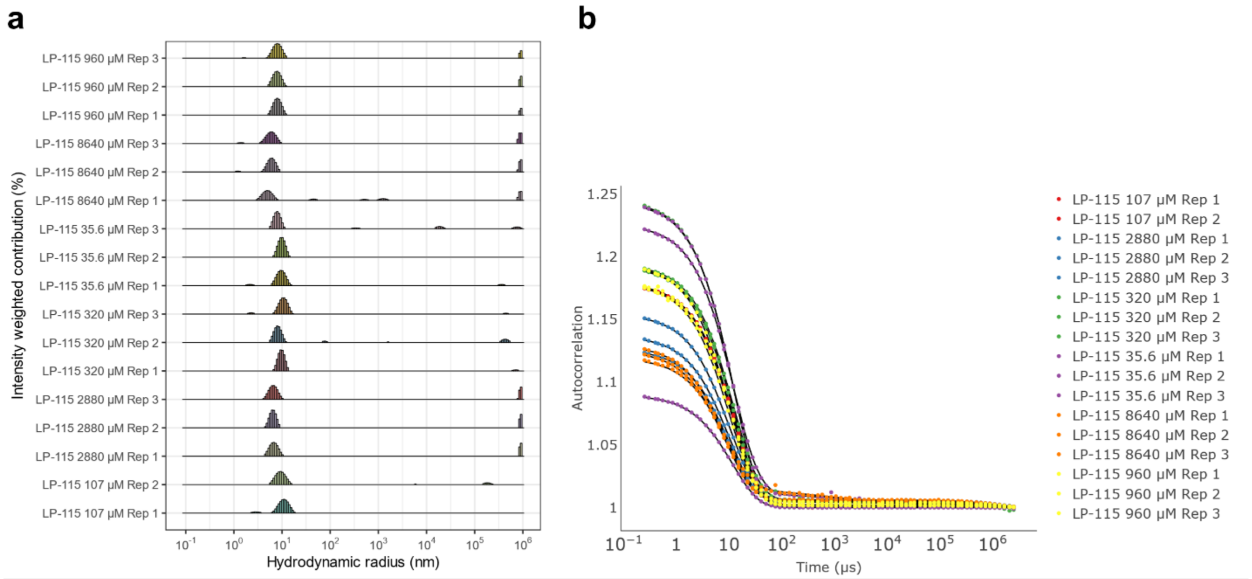


**Fig. S6 | DLS raw data of the AcrABZ-TolC efflux pump with different concentrations of LP-115. a** Derived histograms of the autocorrelation curves (**b**) of the AcrABZ-TolC efflux pump with different concentrations of LP-115 measured by DLS. Each curve presents a mean of ten acquisitions of the same capillary.

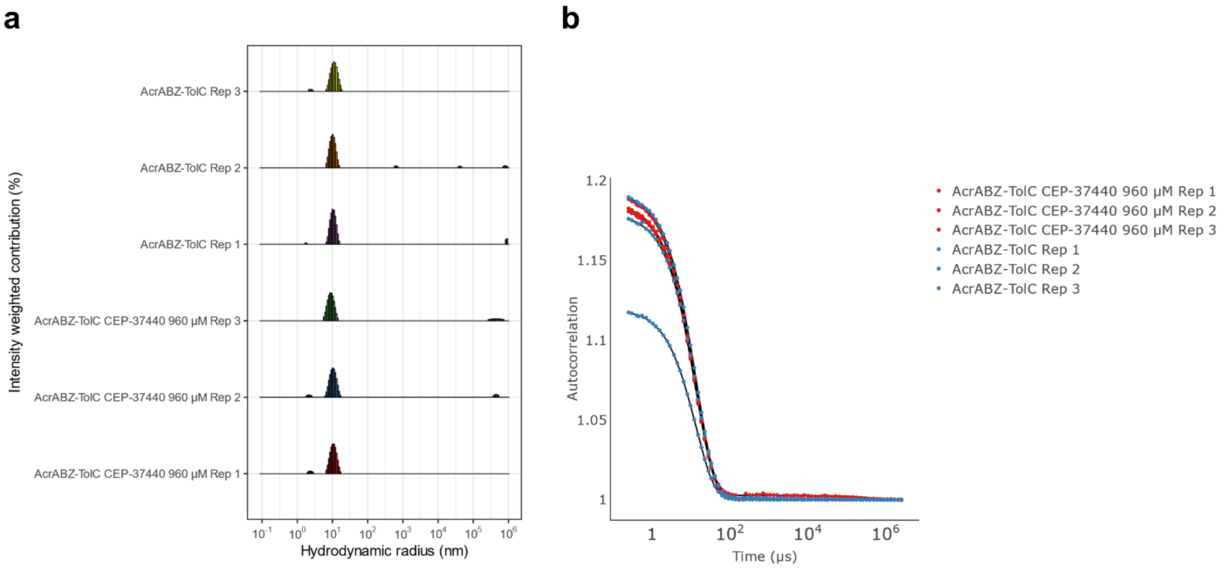


**Fig. S7 | DLS raw data of the AcrABZ-TolC efflux pump in combination with CEP-37440. a** Derived histograms of the autocorrelation curves (**b**) of the AcrABZ-TolC efflux pump with and without CEP-37440 measured by DLS. Each curve presents a mean of ten acquisitions of the same capillary.

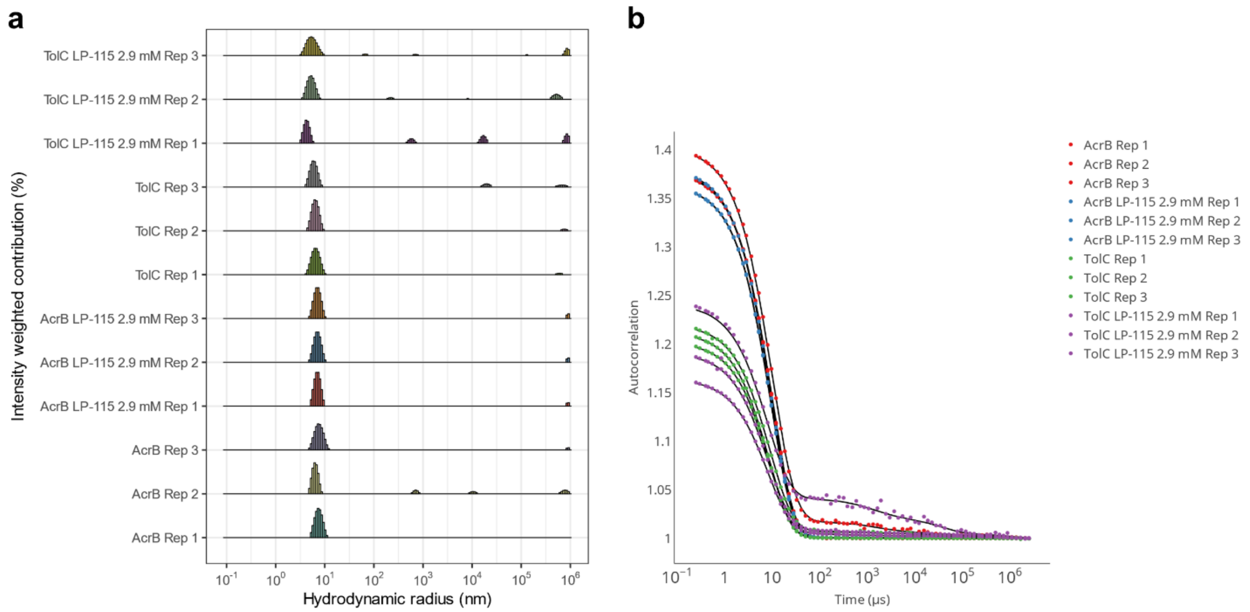


**Fig. S8 | DLS raw data of the efflux pump subunits AcrB and TolC in combination with LP-115. a** Derived histograms of the autocorrelation curves (**b**) of AcrB and TolC with and without LP-115 measured by DLS. Each curve presents a mean of ten acquisitions of the same capillary.

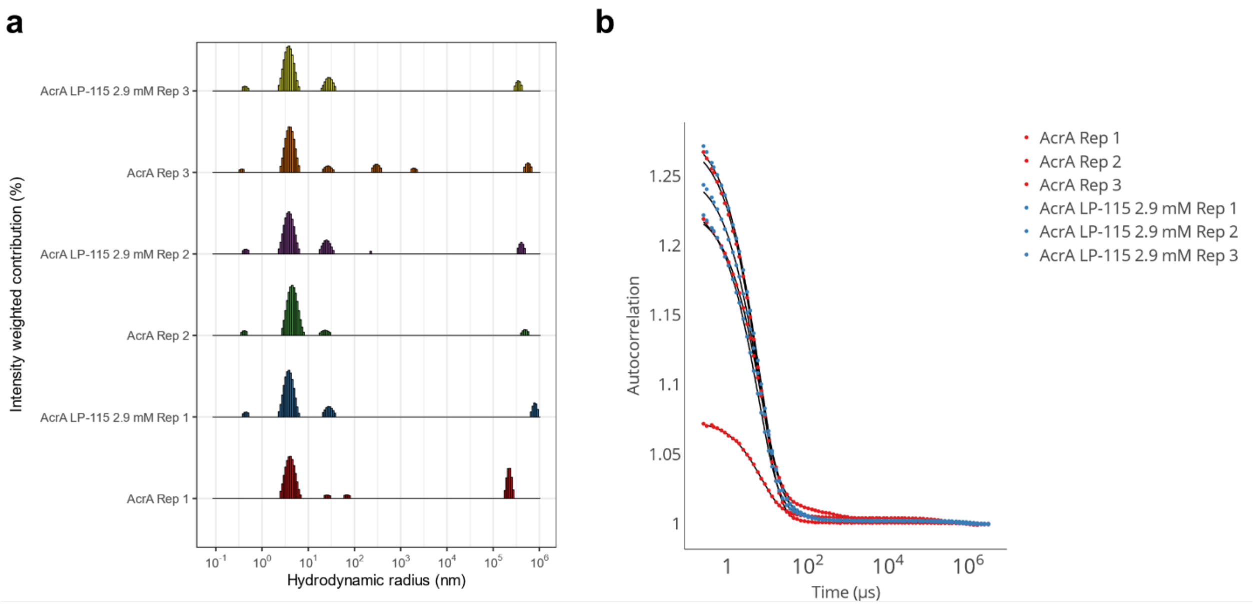


**Fig. S9 | DLS raw data of the efflux pump subunit AcrA in combination with LP-115.** **a** Derived histograms of the autocorrelation curves (**b**) of AcrA with and without LP-115 measured by DLS. Each curve presents a mean of ten acquisitions of the same capillary.

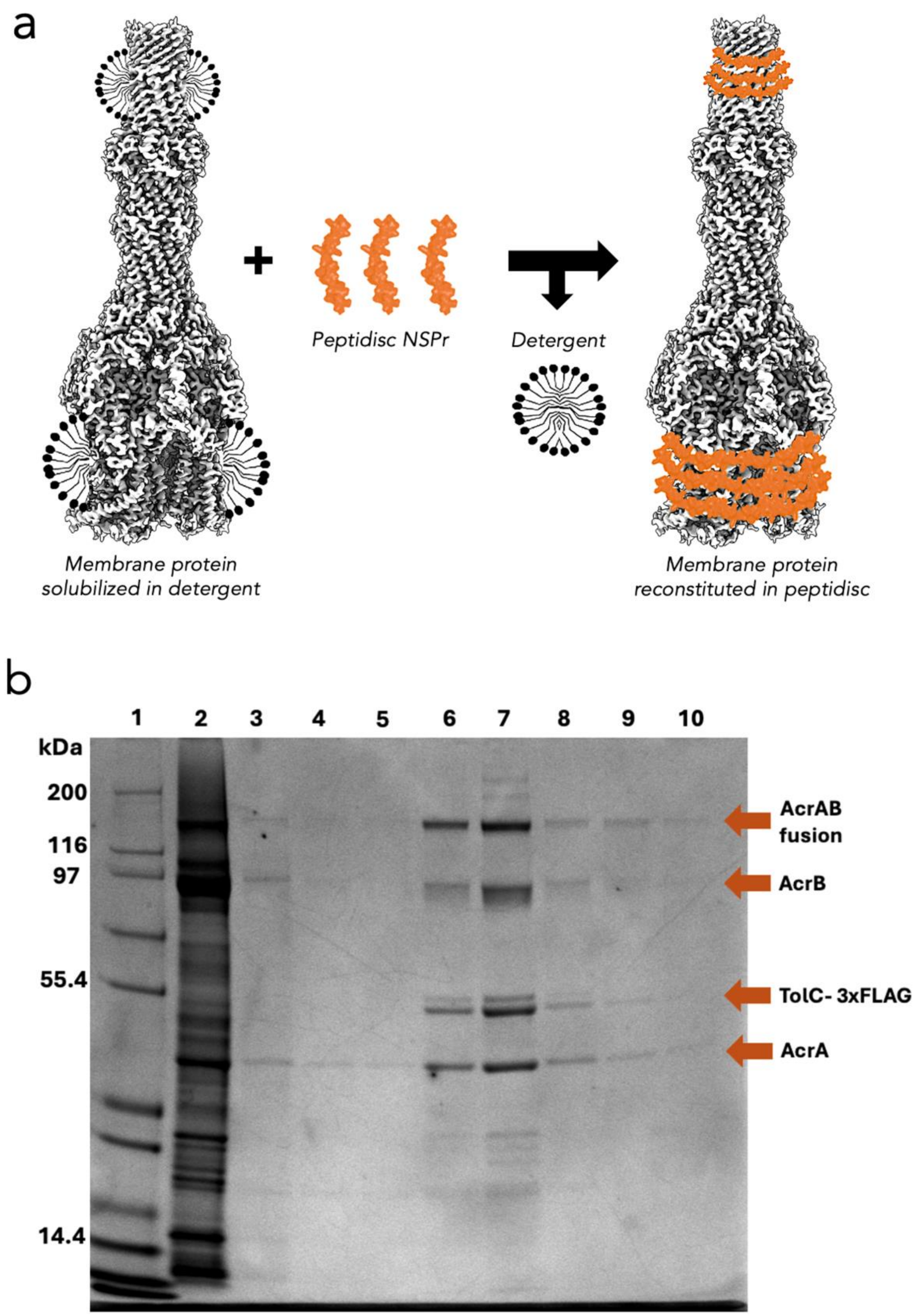


**Fig. S10 | Purification and reconstitution in peptidisc of AcrABZ-TolC. a** Cartoon schematic showing the principle of peptidisc reconstitution. **b** SDS-PAGE gel image of the anti-FLAG beads purification and "on beads" reconstitution results showing all the pump components in the elution fractions after peptidisc reconstitution. Lane 1 Broad-Range SDS-PAGE Standards (Bio-Rad); Lane 2 flowthrough; Lanes 3-5 washes; Lanes 6-10 elution fractions.

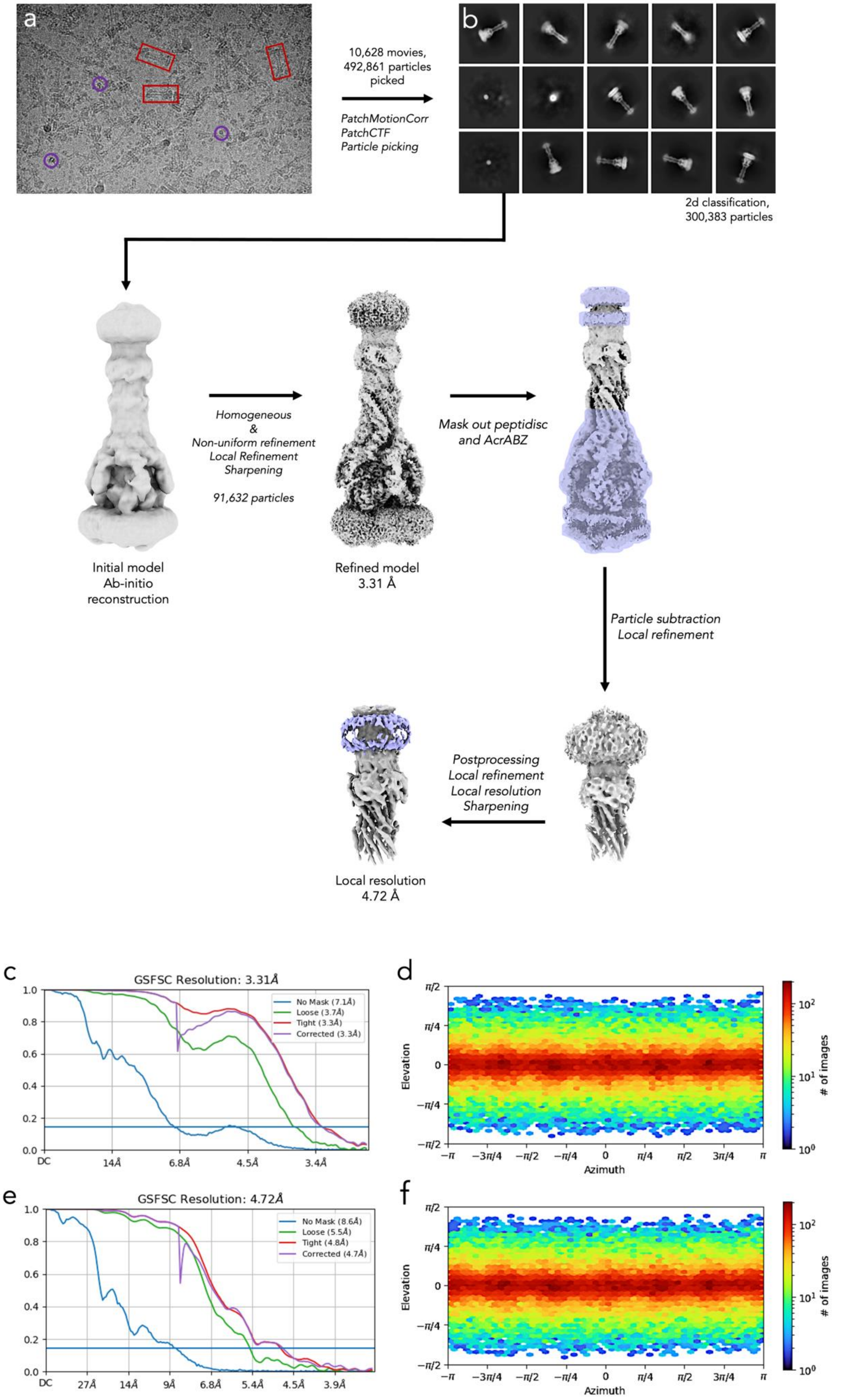
a
b
10,628 movies,
492,861 particles
picked
PatchMotionCorr
PatchCTF
Particle picking
2d classification,
300,383 particles
Homogeneous
&
Non-uniform refinement
Local Refinement
Sharpening
91,632 particles
Initial model
Ab-initio
reconstruction
Refined model
3.31 Å
Mask out peptidisc
and AcrABZ
Particle subtraction
Local refinement
Postprocessing
Local refinement
Local resolution
Sharpening
Local resolution
4.72 Å
c
GSFSC Resolution: 3.31Å
No Mask (7.1Å)
Loose (3.7Å)
Tight (3.3Å)
Corrected (3.3Å)
DC
14Å
6.8Å
4.5Å
3.4Å
d
Elevation
Azimuth
### of images
e
GSFSC Resolution: 4.72Å
No Mask (8.6Å)
Loose (5.5Å)
Tight (4.8Å)
Corrected (4.7Å)
DC
27Å
14Å
9Å
6.8Å
5.4Å
4.5Å
3.9Å
f
Elevation
Azimuth
### of images

**Fig. S11 | Cryo-EM data processing of AcrABZ-TolC in peptidisc bound to LP-115.** **a** A Representative micrograph showing needle-like particles of the reconstituted efflux pump (red squares) and of the TolC-3xFLAG from the top view (purple circles). Single-particle cryo-EM data were processed using cryoSPARC. After motion correction, CTF estimation, automated blob picking and template picking, three rounds of 2D classification were performed to remove bad particles. An ab initio model was built and used to generate a de novo model further refined in C1 using homogeneous refinement and non-uniform refinement, leading to a 3.43 Å resolution map. **b** Representative 2D average classes. **c** Fourier Shell Correlation between half maps. The resolution at 0.143 FSC is indicated by the intersection of the blue line. **d** Angular distribution plot of the particles calculated by cryoSPARC. **e** Fourier Shell Correlation of locally refined map. **f** Angular distribution plot of the particles in local refinement calculated by cryoSPARC.

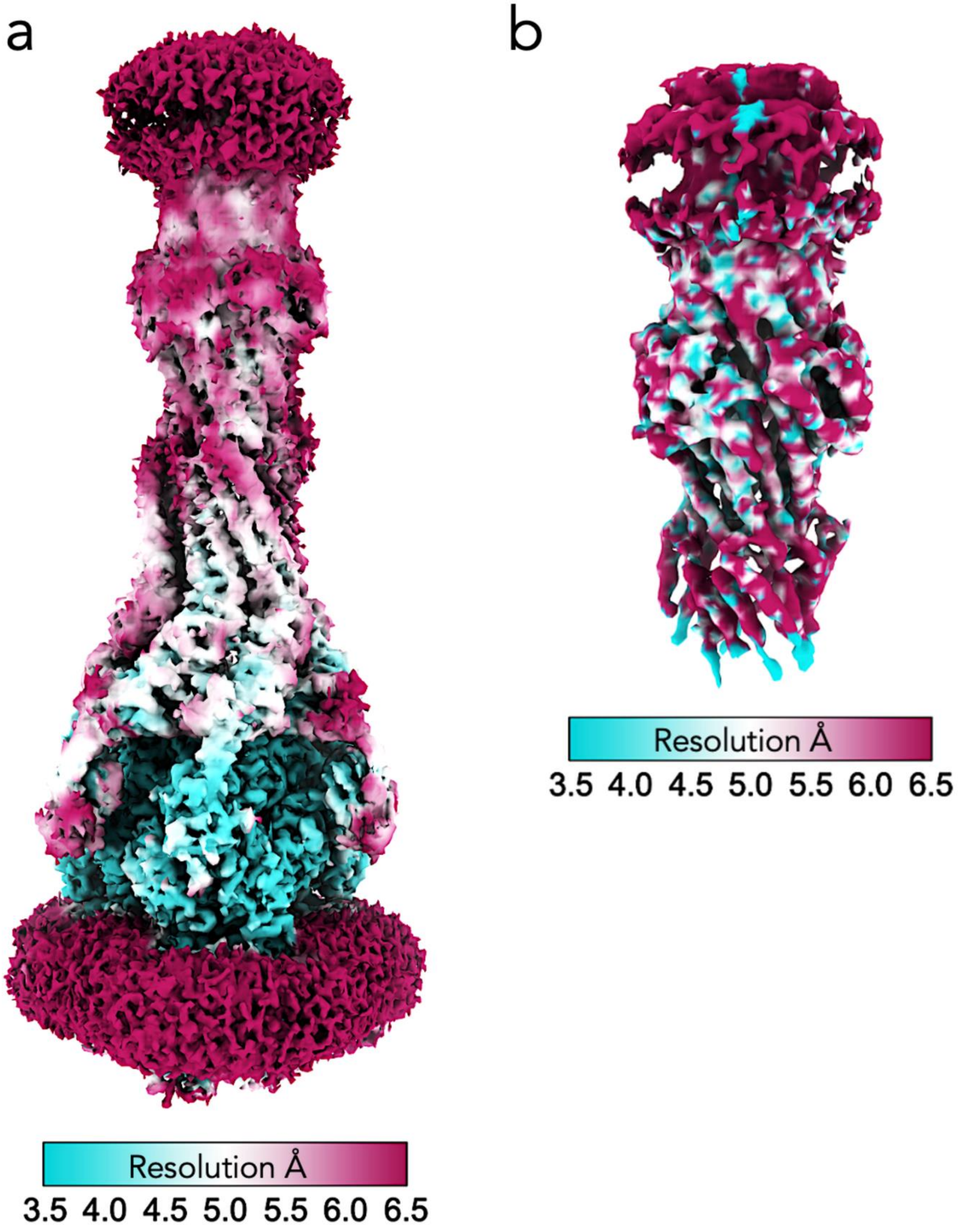


**Fig. S12 | Local resolution analysis of cryo-EM maps. (a).** EM volume map colored according to the local resolution for the final 3D reconstruction of AcrABZ-TolC. **(b).** EM volume map colored according to the local resolution using locres in Cryosparc for the subtracted TolC-AcrA reconstruction where LP115 is bound.

#### Table S3 | Data collection, processing, and model building

| | **AcrABZ-TolC-LP115 pump**<br>PDB ID: XXXX<br>EMDB ID: EMD-XXXXX | **AcrA-TolC-LP115 binary complex pump**<br>PDB ID: XXXX<br>EMDB ID: EMD-XXXXX |
|---|---|---|
| **Data collection and processing** | | |
| Magnification | 130,000x | 130,000x |
| Voltage (kV) | 300 | 300 |
| Electron exposure ($e^-/Å^2$) | 51.49 | 51.49 |
| Defocus range (µm) | -1, -2.5 | -1, -2.5 |
| Pixel size (Å) | 0.652 (binned 2X in EPU) | 0.652 (binned 2X in EPU) |
| Symmetry imposed | C1 | C1 |
| Initial particle images (no.) | 492,861 | 492,861 |
| Final particle images (no.) | 91,632 | 91,632 |
| Map resolution (Å, FSC = 0.143) | 3.31 | 4.72 |
| Particle box size | 540 | 540 |
| **Refinement** | | |
| **Model composition** | | |
| Chains | 15 | 9 |
| Atoms | 50,833 | 12,217 |
| Protein residues | 6701 | 3914 |
| **Bonds (RMSD)** | | |
| Bond lengths (Å) | 0.005 (4) | 0.015 (2) |
| Bond angles (°) | 0.770 (25) | 2.431 (87) |
| **Validation** | | |
| MolProbity score | 2.20 | 3.94 |
| **Ramachandran plot** | | |
| Favored (%) | 92.85 | 89.92 |
| Allowed (%) | 7.59 | 7.46 |
| Outliers (%) | 0.10 | 2.61 |

**Table S4 | Used primers for PCR reaction.**

| Primer | Sequence (5′>3′) |
|---|---|
| pEPF1w-TolC-for | CGCGGGCATATGAAGAAATTGCTCCCCATTCTTATC |
| pEPF1w-TolC-rev | CGCGGGCCTAGGTTATCAGTTACGGAAAGGGTTATGAC |
| pEPFeFt-3xFLAG-for | CGATATTGATTATAAAGATGATGATGATAAAGCTGGTACCCAGTATGACGATAG |
| pEPFeFt-3xFLAG-rev | TTTATCATCATCATCTTTATAATCAATATCGTGGTCTTTATAGTCGCCATCGTGATCTTTGTAATCGGCACCACGGGTTTTCGAAC |
| AcrAS273C_F | GATCAGACCACTGGGTGTATCACCCTACGCGCTATCTTC |
| AcrAS273C_R | GAAGATAGCGCGTAGGGTGATACACCCAGTGGTCTGATC |
| AcrBS258C_F | GTGAATCAGGATGGTTGTCGCGTGCTGCTGCGTGAC |
| AcrBS258C_R | GTCACGCAGCAGCACGCGACAACCATCCTGATTCAC |

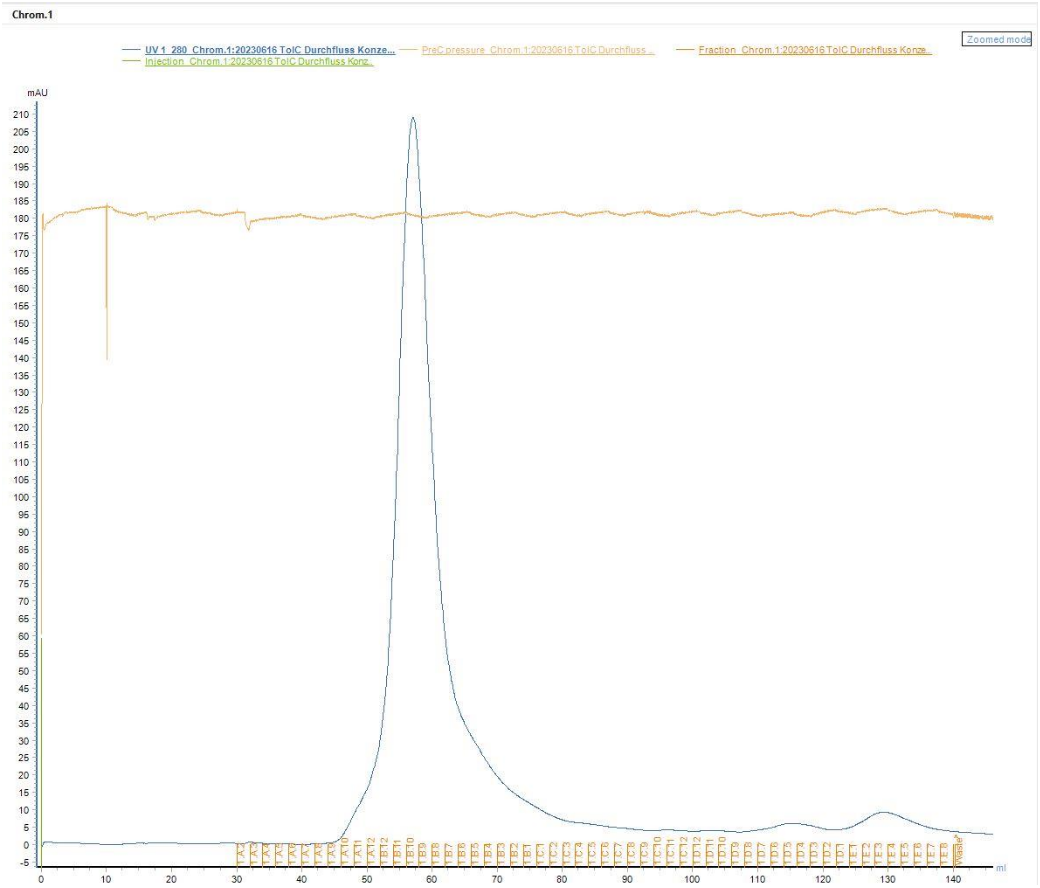


**Fig. S13 | Chromatogram of the size exclusion chromatography (SEC) used to purify TolC.** The chromatogram shows the monitored UV absorption (280 nm) of the sample during the SEC run. It was generated by the ÄKTA surveillance software UNICORN™ 7 (Cytiva). y-axis represents the absorption in mAU, while the x-axis represents the elution volume and the collected fractions.

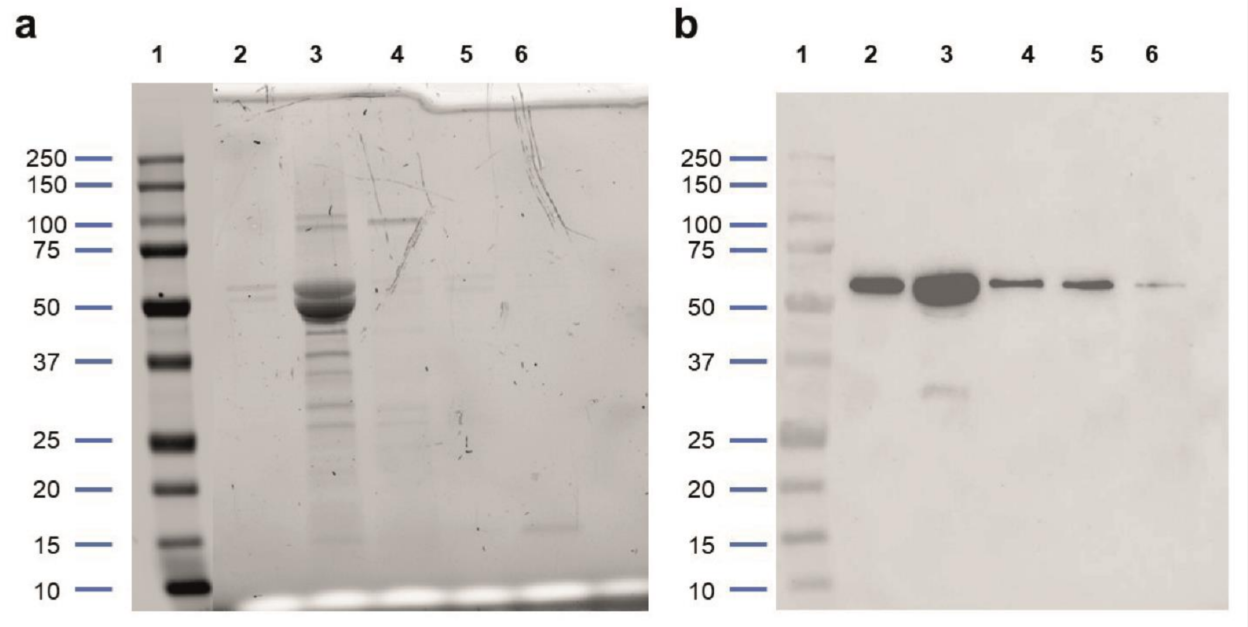


**Fig. S14 | SDS-PAGE and Western Blot documenting the size exclusion chromatography (SEC) step of the TolC purification. a** SDS-PAGE (12 % TGX Stain-Free™ Gel, Bio-Rad Laboratories, Inc.) with different elution fractions along the main elution peak of the SEC step. **b** Anti-His-Antibody developed Western Blot (Anti-His tag Antibody (HRP), mouse monoclonal IgG (His-probe AD1.1.10): sc-53073 (Santa Cruz Biotechnology, Inc.)) with same elution fractions as shown in a. **a**, **b** Lanes were filled with 10 µL of the fraction with loading buffer (4:1) in this order: 1 Precision Plus Protein™ All Blue Prestained Protein Standards (Bio-Rad), 2 SEC Fraction A11, 3 SEC Fraction B10, 4 SEC Fraction B3, 5 SEC Fraction D4, 6 SEC Fraction E3.

**Mass spectrometric measurements – Methods**

Top-down sequencing of the protein

Top-down sequencing for confirmation of the N or C terminus of the protein was performed on a rapifleX MALDI-TOF/TOF instrument (Bruker Daltonics) with a MALDI-ISD (in-source decay) method. The mass range was m/z 1000 to 10000 in positive ionization mode. sDHB (90:10 mixture of 2,5-Dihydroxybenzoic acid and 2-Hydroxy-5-methoxybenzoic acid, Bruker Daltonics) 50 mg/mL 50:50 [v/v] acetonitrile : 0.1% TFA in water was used as a matrix. Data aquisition was performed in FlexControl, for data analysis, FlexAnalysis and Biopharma Compass (each Bruker Daltonics) were used.

Intact mass determination via LC-HRMS

Intact mass determination was performed on a maXis II ETD QTOF mass spectrometer equipped with an Elute UPLC system (both Bruker Daltonics). Desalting was performed using a Thermo ProSwift RP-4H columns (50x1mm) at 50 °C column oven temperature. Solvents used were (A) water + 0.1 % formic acid and (B) acetonitrile + 0.1 % formic acid at a flow rate of 0.3 ml/min, starting at 5 % solvent B for 0.7 min. The gradient went to 15 % B at 1 min runtime, 55 % B at 2.9 min, 90 % B at 3 min, followed by reequilibration to 5 % B. The whole method has a runtime of 5 min. The MS measurement was performed in positive ionization mode in a mass range of m/z 400 – 2500. Data aquisition was performed using HiStar and oTOF control (each Bruker Daltonics), for data analysis DataAnalysis (Bruker Daltonics) was used. The protein mass was determined using the Maximum Entropy algorithm for deconvolution in DataAnalysis.

In-gel digestion

In-gel digestion was done as previously described[1]. Shrinking and swelling was performed with 100 % ACN and 100 mM NH4HCO3. In-gel reduction was achieved with 10 mM dithiothreitol (dissolved in 100 mM NH4HCO3). Alkylation was performed with 55 mM iodacetamide (dissolved in 100 mM NH4HCO3). Proteins in the gel pieces were digested by covering them with a trypsin solution (8 ng/µL sequencing-grade trypsin, dissolved in 50 mM NH4HCO3) and incubating the mixture at 37 C for overnight. Tryptic peptides were yielded by extraction with 2% FA, 90% ACN. The extract was evaporated. For LC-MS/MS analysis, samples were dissolved in 20 µL 0.1 % FA.

LC-MS/MS

Chromatographic separation of peptides was achieved with a two-buffer system (buffer A: 0.1% FA in H2O, buffer B: 0.1% FA in ACN) on a nano-UPLC (Dionex Ultimate 3000 UPLC system, Thermo Fisher). Attached to the UHPLC was a peptide trap (100 µm x 20 mm, 100 Å pore size, 5 µm particle size, C18, Nano Viper, Thermo Fisher) for online desalting and purification, followed by a 25 cm C18 reversed-phase column (75 µm x 250 mm, 130 Å pore size, 1.7 µm particle size, peptide BEH C18, nanoEase, Waters). Peptides were separated using an 80 min method with linearly increasing ACN concentration from 2% to 30% ACN over 60 minutes.

MS/MS measurements were performed on a quadrupole-orbitrap hybrid mass spectrometer (QExactive, Thermo Fisher Scientific). Eluting peptides were ionized using a nano-electrospray ionization source (nano-ESI) with a spray voltage of 1,800 and analyzed in data dependent acquisition (DDA) mode. For each MS1 scan, ions were accumulated for a maximum of 240 milliseconds or until a charge density of 1 x 106 ions (AGC Target) was reached. Fourier-transformation based mass analysis of the data from the orbitrap mass analyzer was performed covering a mass range of 400 – 1,200 m/z with a resolution of 70,000 at m/z = 200. Peptides being responsible for the 15 highest signal intensities per precursor scan with a minimum AGC target of 5 x 103 and charge state from +2 to +5 were isolated within a 2 m/z isolation window and fragmented with a normalized collision energy of 25% using higher energy collisional dissociation (HCD). MS2 scanning was performed, covering a mass range starting at 100 m/z and accumulated for 50 ms or to an AGC target of 1 x 105 at a resolution of 17,500 at m/z = 200. Already fragmented peptides were excluded for 20 s.

Data analysis

LC-MS/MS data were searched with the Sequest algorithm integrated into the Proteome Discoverer software (v2.41.15, Thermo Fisher Scientific) against a reviewed Thermoplasma acidophilum Swissprot database, obtained in September 2023, containing 1,543 entries plus the sequence of the protein of interest. Carbamidomethylation was set as a fixed modification for cysteine residues. The oxidation of methionine, and pyro-glutamate formation at glutamine residues at the peptide N-terminus, as well as acetylation of the protein N-terminus were allowed as variable modifications. A maximum number of two missing tryptic cleavages was set. Peptides between 6 and 144 amino acids were considered. A strict cutoff (FDR < 0.01) was set for peptide and protein identification. Quantification was performed using the Minora Algorithm, implemented in Proteome Discoverer. Obtained protein abundances were log2-transformed and normalized by column-median normalization.

#### Mass spectrometric measurements – Results and Discussion

TolC appeared on the SDS-PAGE (Fig. S14) as two bands at the height of the 50 kDa marker protein. This has been observed by others and explained with a proteolytic cleavage of TolC at Arg459[2]. To prove this and further check the effect that storage of TolC at 4 °C during the biophysical measurements had, the Protein was analysed extensively by the Core Facility Mass Spectrometric Proteomics at the Medical Center Hamburg-Eppendorf (UKE). Both bands were digested with Trypsin and identified in LC-MS/MS measurements as TolC (Fig. S11b). However, the masses measured in LC-HRMS were smaller than expected, approximately 48 kDa (Fig. S11a) instead of 51.5 kDa. N-terminal sequencing proved the N-terminus to be complete (Fig. S11b), starting with the first amino acid after the signal peptide. The cleavage of the signal peptide proves that TolC is correctly integrated in the outer membrane during protein expression. A C-terminal truncation from amino acids Q437 to A439 is potentially responsible for the decreased size, since the C-terminal deletion of these amino acids results in 47.9 to 48.1 kDa sized TolC derivatives (analyzed with the Expasy ProtParam tool (SIB Swiss Institute of Bioinformatics)). For the *in-silico* docking studies, X-ray structures have been used that lack the amino acids after E428 (as usual for published structures of TolC), so docking to the truncated amino acids of TolC after Q437 was not possible and it is not expected to be part of the binding site. The amino acids after Q437 form the flexible periplasmic chain of TolC and given that the binding site is expected at the periplasmic tip of TolC, the truncation presumably does not affect the binding.

**a**

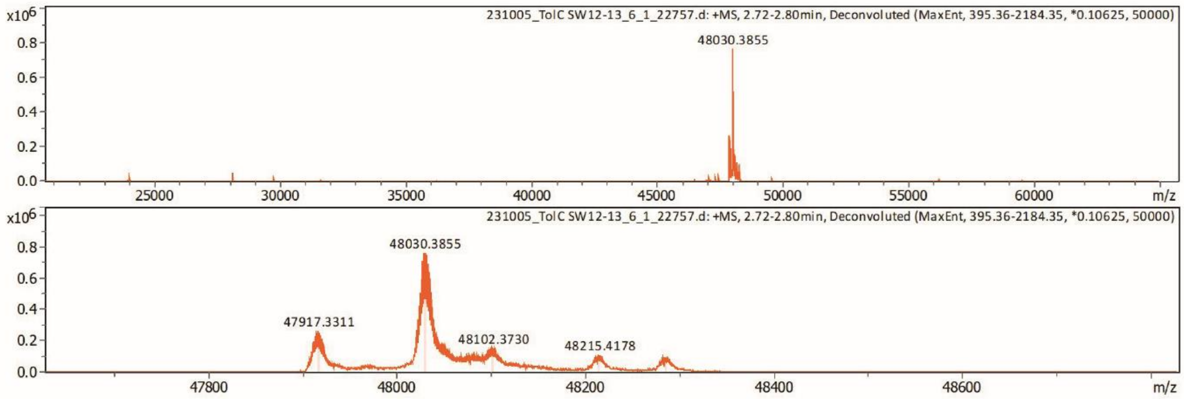


**b**

**ENLMQVYQQA** **RLSNPELR**KS **AADRDAAFEK** **INEARSP**LLP QLGLGAD**Y**TY SNGYR**DANGI** **NSNATSASLQ** **LTQSIFDMSK**

WR**ALTLQEK**A AGIQDVTYQT DQQTLILNTA TAYFNVLNAI DVLSYTQAQK **EAIYRQLDQT** **TQRFNVGLVA** **ITDVQNARAQ**

**YDTVLANEVT** **ARNNLDNAVE** **QLRQITGNYY** **PELAALNVEN** **FKTDKPQPVN** **ALLKEAEKRN** **LSLLQARLSQ** **DLAREQIRQA**

**QDGHLPTLDL** **TASTGISDTS** **YSGSK**TR**GAA** **GTQYDDSNMG** **QNKVGLSFSL** **PIYQGGMVNS** **QVKQAQYNFV** **GASEQLESAH**

**R**SVVQTVR**SS** **FNNINASISS** **INAYKQAVVS** **AQSSLDAMEA** **GYSVGTRTIV** **DVLDATTTLY** **NAKQELANAR** **YNYLINQLNI**

**K**SALGTLNEQ DLLALNNALS KPVSTNPENV APQTPEQNAI ADGYAPDSPA PVVQQTSART TTSNGHNPFR NHHHHLEHHH HHH

**Fig. S15 | Mass spectrometric analysis of the TolC batch after the biophysical measurements. a** LC-HRMS measurements with a main peak at 48.0 kDa and two smaller ones at 47.9 and 48.1 kDa. **b** TolC sequence without the signal peptide. Amino acids detected with MALDI-ISD method or LC-MS/MS (after Trypsin digestion) are presented in bold.

#### Nano Differential Scanning Fluorimetry (nDSF) – Method

The nDSF measurements were conducted with Tycho NT.6 (NanoTemper Technologies). The protein solution was either measured undiluted or, for the DMSO tolerance, diluted in the respective buffer with 5 or 25 % DMSO to a final protein concentration of 5 µM. A dilution series of DMSO in PBS pH 7.4 and 0.03% DDM was prepared, and protein was added. After an incubation of 15 minutes, Tycho capillaries (NanoTemper Technologies) were loaded with the respective samples and inserted into the instrument for measurement. After a visual inspection of the melting curves, the determined melting temperatures were compared.

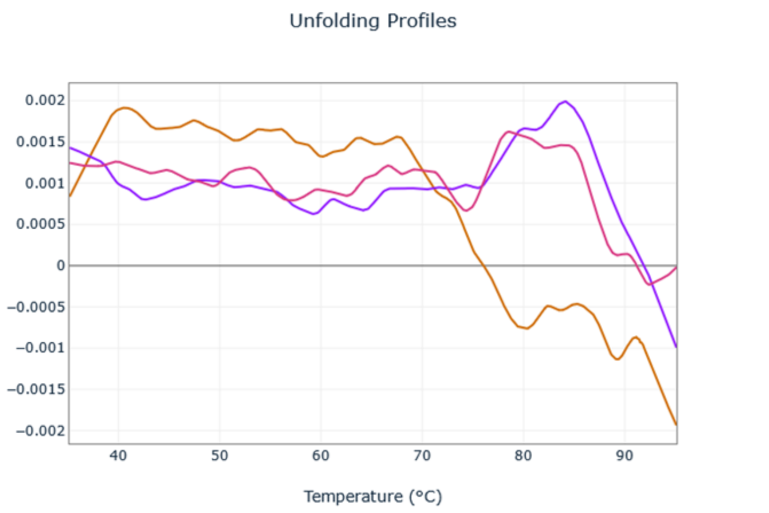


**Fig. S16 | nDSF measurements of TolC with different DMSO concentrations.** First derivative of the melting curve of 5 µM TolC with 5% (pink) and 25% DMSO (orange), in comparison with the melting curve of 5 µM TolC without addition of DMSO (violet).

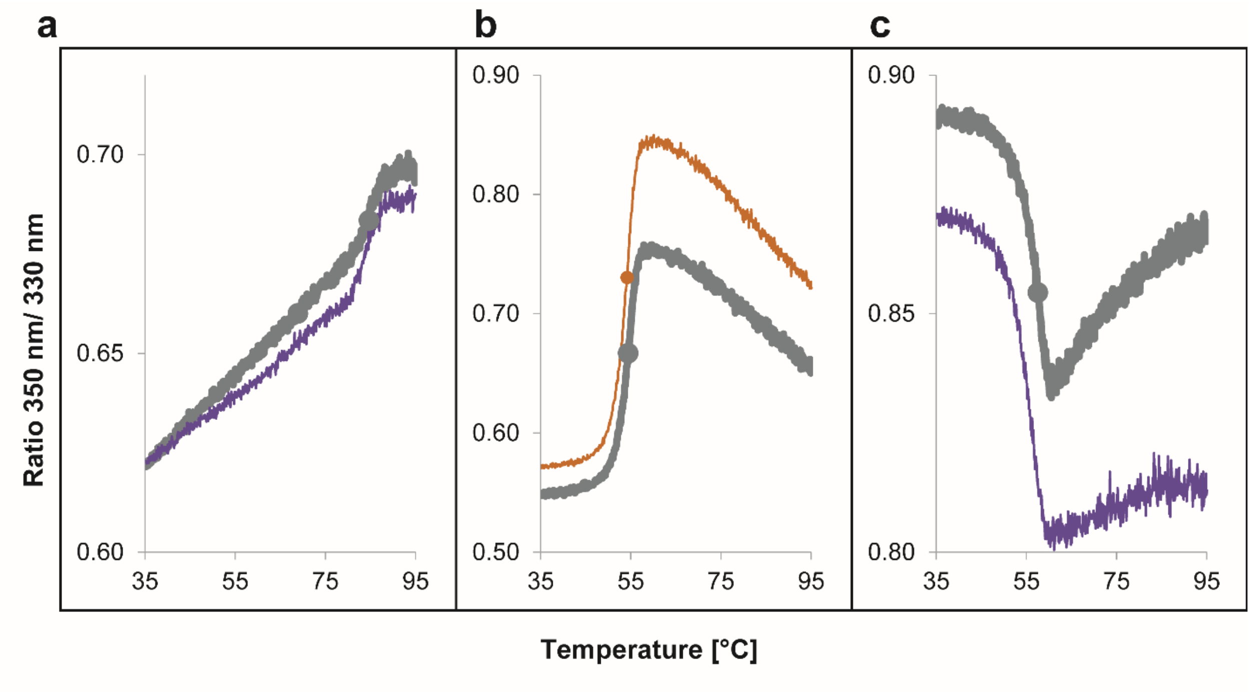


**Fig. S17 | nDSF measurements to ensure protein stability after labeling.** Melting curves of the proteins before (gray) and after labeling (TolC: **a**, violet; AcrA: **b**, orange; AcrB: **c**, violet).

**Table S5 | Degree of Labeling of the labeled proteins.** Absorption at 280 and 650 nm was measured using a NanoDrop® ND-1000 UV-Vis Spectrophotometer (Thermo Scientific GmbH) at 280 and 650 nm. The degree of labeling was calculated following the manufacturer's (NanoTemper Technologies) instructions for the labeling kit used.

| Protein | Degree of labeling |
|---|---|
| AcrA | 0.91 |
| AcrB | 0.48 |
| TolC | 0.40 |

#### Synthesis of CEP-37440 fragments

**General remarks.** All chemicals, reagents and solvents were purchased from commercial sources and were used without further purification. Synthesis of compounds **2** to **6**, **LP-115** was performed following modified literature procedures[1]. Fragment 2-amino-N-methylbenzamide, 95% (**OSM-2**) was purchased from Enamine Ltd (cat.nr.: EN300-17933) and used without further purification. Thin layer chromatography (TLC) was performed on silica gel using Merck TLC Silca gel 60 $F_{254}$ Aluminium sheets and was visualized by UV lamp. The direct phase flash column chromatography was carried out using Kieselgel silica gel (35 -70 microm). The reverse phase silica gel chromatography was performed using Biotage purification system with prepacked columns. NMR spectra were recorded on 400 MHz Bruker spectrometer with chemical shift values (δ) in parts per million using the residual solvent signal as an internal standard. The multiplicities are denoted as follows: s, singlet; d, doublet; dd, doublet of doublets; t, triplet; qd, quartet of doublet; m, multiplet; br s, broad singlet. HRMS analyses were performed on a hybrid QToF spectrometer equipped with an electrospray ion source. The purity of all the tested compounds was estimated by HPLC (Empower 3 or SHIMADZU LabSolutions, UV detection at $\lambda$ = 210 nm) analysis on an Adamas C18 column size: 4.6 x 150 mm, mobile phase: MeCN - 0.1% $H_3PO_4$, flow rate: 1.0 ml/ min, detector: UV 210 nm and UV 254 nm column temperature: 40°C, sample concentration: 0.5 mg/ml. Abbreviations: HFIP - hexafluoro-2-propanol. PGME - 1-methoxy-2-propanol.

[1] Allwein, S. P.; Mowrey, D. R.; Petrillo, D. E.; Reif, J. J.; Purohit, V. C.; Milkiewicz, K. L.; Gilmartin, G. J. Development of a Process Route to the FAK/ALK Dual Inhibitor TEV-37440. *Organic Process Research & Development*. **2017,** 21(5), 740–747. DOI:10.1021/acs.oprd.7b00070.

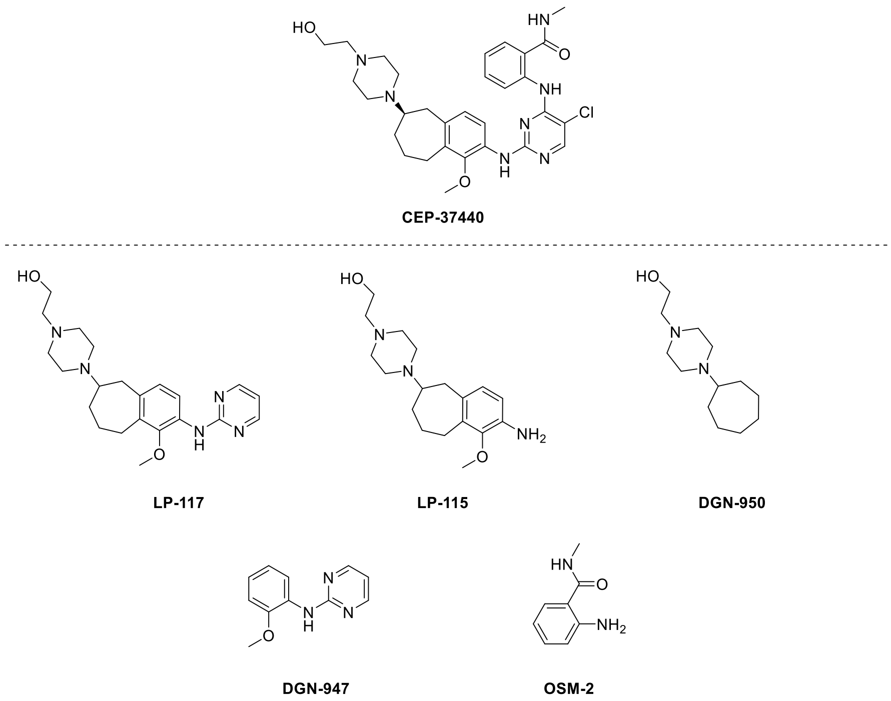


**Figure S18 | Deconstruction of CEP-37440 into fragments.**

#### Scheme S1

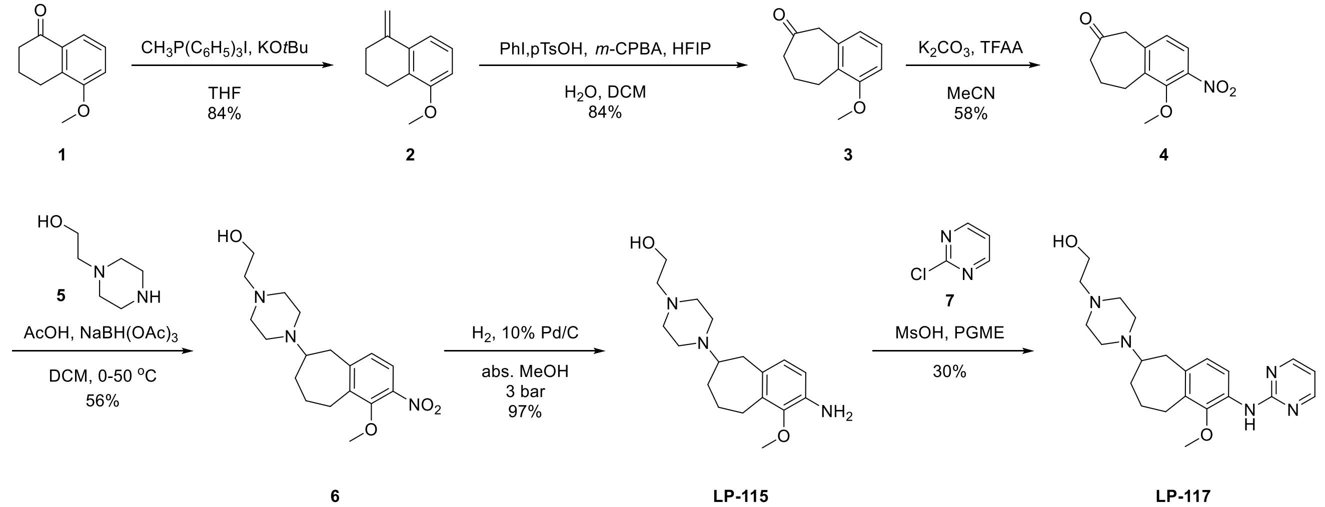


**5-Methoxy-1-methylene-1,2,3,4-tetrahydronaphthalene (2).** To a suspension of 5-methoxy-3,4-dihydronaphthalen-1(2H)-one (**1**) (1.23 g, 6.98 mmol) and $CH_3P(C_6H_5)_3I$ (3.67 g, 9.07 mmol) in dry THF (20 ml) *t*-BuOK (1.49 g, 13.26 mmol) dissolved in dry THF (30 ml) was added dropwise within 30 minutes. The resulted mixture was stirred at r.t. overnight, concentrated in vacuum and dissolved in EtOAc (70 ml), washed with $H_2O$ (4 x 20 ml) and brine (20 ml), dried over $Na_2SO_4$ and evaporated. The crude product was purified by silica gel column chromatography (PET/EtOAc 30:1 v/v) to obtain compound **2** as a yellow oil (1.02 g, 84%). $^1$H NMR (400 MHz, $CDCl_3$) δ 7.29 (d, *J* = 7.8 Hz, 1H), 7.15 (t, *J* = 8.0 Hz, 1H), 6.76 (d, *J* = 7.9 Hz, 1H), 5.48 (s, 1H), 4.98 (s, 1H), 3.84 (s, 3H), 2.77 (t, *J* = 6.4 Hz, 2H), 2.54 – 2.48 (m, 2H), 1.94 – 1.86 (m, 2H) ppm.

**1-Methoxy-5,7,8,9-tetrahydro-6H-benzo[7]annulen-6-one (3).** Iodobenzene (0.19 ml, 1.66 mmol), 70% *m*-CPBA (0.41 g, 1.66 mmol) and *p*-toluenesulfonic acid monohydrate (0.32 g, 1.66 mmol) were dissolved in dry DCM (10 ml). HFIP (1.7 ml) was added and the mixture was stirred at r.t. for 45 minutes. The solution was cooled to 0 $^oC$ (ice-bath) with further addition of $H_2O$ (0.68 ml) and 5-methoxy-1-methylene-1,2,3,4-tetrahydronaphthalene (**2**) (0.25 g, 1.44 mmol), and stirred at 0 $^oC$ (ice-bath) for 2 hours. The resulted mixture was diluted with DCM (20 ml) and saturated aqueous $NaHCO_3$ (10 mL) was added to pH ~8. The organic layer was washed with $H_2O$ (3 x 10 ml) and brine (10 ml), dried over $Na_2SO_4$ and evaporated. The crude product was purified by silica gel column chromatography (PET/EtOAc 10:1 v/v) to obtain compound **3** as a yellow oil (0.23 g, 84%). $^1$H NMR (400 MHz, $CDCl_3$) δ 7.14 (t, 1H), 6.82 (d, *J* = 7.8 Hz, 1H), 6.77 (d, *J* = 7.5 Hz, 1H), 3.82 (s, 3H), 3.71 (s, 2H), 3.05 – 2.97 (m, 2H), 2.53 (t, *J* = 6.9 Hz, 2H), 2.00 – 1.92 (m, 2H) ppm.

**1-Methoxy-2-nitro-5,7,8,9-tetrahydro-6H-benzo[7]annulen-6-one (4).** To $KNO_3$ (0.41 g, 4.03 mmol) in MeCN (1.5 ml) and trifluoroacetic anhydride (3 ml) at 0 $^oC$ (ice-bath) 1-methoxy-5,7,8,9-tetrahydro-6H-benzo[7]annulen-6-one (**3**) (0.75 g, 3.95 mmol) in MeCN (5 ml) was added dropwise. The reaction mixture was stirred for 2.5 hours while warming to r.t. MeOH (3 ml) was added and stirred briefly at 0 $^oC$ (ice-bath). The reaction mixture was concentrated in vacuum and dissolved in DCM (50 ml), worked-up with sat. $NaHCO_3$ to pH~7. The organic layer was washed with $H_2O$ (20 ml) and brine (25 ml), dried over $Na_2SO_4$. The crude product was purified by silica gel column chromatography (PET/EtOAc gradient from 10:1 to 10:2 v/v, the desired *ortho* isomer was eluting later) to obtain compound **4** (0.54 g, 58% with ~10% *para* isomer). $^1$H NMR (400 MHz, $CDCl_3$) δ 7.68 (d, *J* = 8.3 Hz, 1H), 7.04 (d, *J* = 8.3 Hz, 1H), 3.90 (s, 3H), 3.78 (s, 2H), 3.11 – 3.07 (m, 2H), 2.58 (t, *J* = 6.9 Hz, 2H), 2.08 – 2.01 (m, 2H) ppm.

**2-(4-(1-Methoxy-2-nitro-6,7,8,9-tetrahydro-5H-benzo[7]annulen-6-yl)piperazin-1-yl)ethan-1-ol (6).** 1-Methoxy-2-nitro-5,7,8,9-tetrahydro-6H-benzo[7]annulen-6-one (**4**) (0.11 g, 0.47 mmol) was dissolved in dry DCM (5 ml) and 2-(piperazin-1-yl)ethan-1-ol (**5**) (0.17 ml, 1.40 mmol) and glacial acetic acid (0.3 ml, 4.70 mmol) were added and the reaction mixture was stirred at 50 $^oC$ for 2 hours and cooled to 0 $^oC$ (ice-bath) before adding $NaBH(OAc)_3$ (0.40 g, 1.88 mmol). The mixture was stirred at r.t. overnight, poured into ice and sat. $NaHCO_3$ solution (5 ml), diluted with EtOAc (40 ml), washed with sat. $NaHCO_3$ solution (5 ml) to pH~8, $H_2O$ (4 x 15 ml) and brine (15 ml), dried over $Na_2SO_4$. The crude product was purified by silica gel column chromatography (DCM/MeOH 10:1 v/v) followed by reverse phase flash column chromatography (BIOTAGE$^{(R)}$ SNAP Cartrige KP-C18-HS, linear gradient elution from 0% to 60% $CH_3CN$) to obtain product **6** (0.45 g, 56%). $^1$H NMR (400 MHz, $CDCl_3$) δ 7.52 (d, *J* = 8.3 Hz, 1H), 6.95 (d, *J* = 8.3 Hz, 1H), 3.78 (s, 3H), 3.62 – 3.58 (m, 2H), 3.26 – 3.18 (m, 1H), 2.96 – 2.78 (m, 3H), 2.71 – 2.65 (m, 2H), 2.60 – 2.52 (m, 8H), 2.41 – 2.34 (m, 2H), 2.08 – 2.01 (m, 2H), 1.84 – 1.70 (m, 1H), 1.34 – 1.23 (m, 1H) ppm.

**2-(4-(2-Amino-1-methoxy-6,7,8,9-tetrahydro-5H-benzo[7]annulen-6-yl)piperazin-1-yl)ethan-1-ol (LP-115).** 2-(4-(1-Methoxy-2-nitro-6,7,8,9-tetrahydro-5H-benzo[7]annulen-6-yl)piperazin-1-yl)ethan-1-ol **6** (1,00 g, 2.86 mmol) was dissolved in abs. MeOH (33 ml) and 10% Pd/C (100 mg) was added. The reaction was hydrogenated for 4 hours at 3 bar, filtrated through fine filter and concentrated (0.89 g, 97%). $^1$H NMR (400 MHz, $CDCl_3$) δ 6.74 (d, *J* = 7.9 Hz, 1H), 6.51 (d, J = 7.9 Hz, 1H), 3.68 (s, 3H), 3.64 – 3.56 (m, 2H), 3.26 – 3.14 (m, 1H), 2.85 – 2.65 (m, 4H), 2.65 – 2.43 (m, 8H), 2.43 – 2.27 (m, 2H), 2.14 – 2.01 (m, 2H), 1.82 – 1.66 (m, 1H), 1.41 – 1.23 (m, 1H) ppm.

**2-(4-(1-Methoxy-2-(pyrimidin-2-ylamino)-6,7,8,9-tetrahydro-5H-benzo[7]annulen-6-yl)piperazin-1-yl)ethan-1-ol (LP-117).** 2-(4-(2-Amino-1-methoxy-6,7,8,9-tetrahydro-5H-benzo[7]annulen-6-yl) piperazin-1-yl) ethan-1-ol (**LP-115**) (80 mg, 0.25 mmol) and 2-chloropyrimidine (**7**) (72 mg, 0.63 mmol) were dissolved in PGME (0.9 ml) and MsOH (0.04 ml, 0.68 mmol) was added. The reaction mixture was heated at 90 $^o$C for 4 hours, and then diluted with DCM (40 ml) and washed with sat. $NaHCO_3$ (3 x 5 ml) solution and brine (10 ml), dried over $Na_2SO_4$ and evaporated. The crude product was purified by silica gel column chromatography (DCM/MeOH gradient from 100:2 to 100:7 v/v) to obtain product **LP-117** as a yellowish solid (30 mg, 30%). $^1$H NMR (400 MHz, $CDCl_3$) δ 8.43 (d, *J* = 4.8 Hz, 2H), 8.18 (d, *J* = 8.3 Hz, 1H), 7.60 (s, 1H), 6.98 (d, *J* = 8.2 Hz, 1H), 6.72 (t, *J* = 4.8 Hz, 1H), 3.74 (m, 2H), 3.72 (s, 3H), 3.27 - 3.18 (m, 1H), 3.05 - 2.79 (m, 10H), 2.75 (m, 2H), 2.57 (br s, 1H), 2.42 (m, 1H), 2.13 (m, 2H), 1.80 (qd, *J* = 12.1, 3.1 Hz, 1H), 1.45 - 1.32 (m, 1H) ppm. $^{13}$C NMR (100 MHz, $CDCl_3$) δ 160.2, 158.1, 146.4, 135.3, 131.5, 125.6, 117.3, 112.8, 112.7, 64.3, 61.6, 60.2, 57.1, 52.3, 46.7, 38.1, 33.5, 26.2, 26.2 ppm. HRMS ESI (m/z): Calculated for $C_{22}H_{32}N_5O_2$ $[M+H]^+$ 398.2556 found 398.2568. Impurities: By HPLC analysis on *Apollo C18*: at 210 nm - 7.44%; at 254 nm - 1.59%.

**Scheme S2**

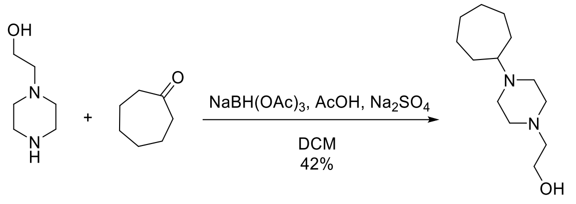


**8** **9** **DGN-950**

**2-(4-Cycloheptylpiperazin-1-yl)ethan-1-ol (DGN-950).** Cycloheptanone (**9**) (0.2 ml, 1.65 mmol) was dissolved in dry DCM (8 ml), 2-(piperazin-1-yl)ethan-1-ol (**8**) (0.2 ml, 1.65 mmol) and glacial acetic acid (0.9 ml, 16.51 mmol) were added. The reaction mixture was stirred at 50 $^o$C for 2 hours and cooled to 0 $^o$C (ice-bath) before adding $NaBH(OAc)_3$ (1.40 g, 6.60 mmol) in portions. The mixture was stirred at r.t. overnight, poured into ice and sat. $NaHCO_3$ solution (15 ml), diluted with EtOAc (60 ml), washed with sat. $NaHCO_3$ solution (10 ml) to pH~8, $H_2O$ (4 x 20 ml) and brine (25 ml), dried over $Na_2SO_4$ and evaporated. The crude product was purified by silica gel column chromatography (DCM/MeOH 10:1 v/v) to obtain product **DGN-950** (158 mg, 42%). $^1$H-NMR (400 MHz, $CDCl_3$) δ 3.61 - 3.58 (m, 2H), 2.58 - 2.51 (m, 11H), 1.85-1.79 (m, 2H), 1.71 - 1.63 (m, 2H),

1.58 - 1.36 (m, 8H) ppm. $^{13}$C-NMR (101 MHz, $CDCl_3$) δ 64.9, 59.6, 57.8, 53.5, 48.3, 29.9, 28.2, 25.8 ppm. GC/MS: 226.2.

**Scheme S3**

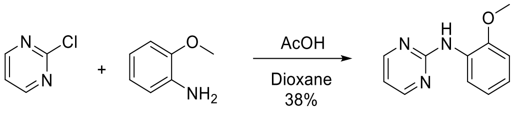


**N-(2-Methoxyphenyl)pyrimidin-2-amine (DGN-947).** 2-Chloropyrimidine (**10**) (400 mg, 3.49 mmol) and 2-methoxyaniline (**11**) (0.5 ml, 4.19 mmol) were dissolved in 1,4-dioxane (7 ml) and acetic acid (1.00 ml, 17.46 mmol) was added. The reaction mixture was stirred at 110 $^{o}$C until complete consumption of starting material **10**. Sat. $NH_4Cl$ aqueous solution (10 ml) was added and the mixture was extracted with DCM (3 x 15 ml), organic layers were combined and dried over $Na_2SO_4$ and evaporated. The crude product was purified by silica gel column chromatography (Hex/EtOAc) to obtain product **DGN-947** (267 mg, 38%). $^{1}$H-NMR (400 MHz, DMSO-$d_6$) δ 8.45 (d, *J* = 4.8 Hz, 2H), 8.13 (dd, *J* = 7.9, 1.5 Hz, 1H), 8.06 (s, 1H), 7.06 - 6.99 (m, 2H), 6.96-6.91 (m, 1H), 6.84 (t, *J* = 4.8 Hz, 1H), 3.85 (s, 3H) ppm. $^{13}$C-NMR (100 MHz, DMSO-$d_6$) δ 159.9, 158.1, 149.1, 128.5, 122.8, 120.3, 120.3, 112.6, 110.8, 55.7 ppm. HRMS (ESI) m/z: Calculated for $C_{11}H_{12}N_3O$ $[M+H]^+$ 202.0980, found 202.0984. Impurities: By HPLC analysis on *Apollo C18*: at 210 nm - 2.15%; at 254 nm - 0.93%

#### $^1$H NMR (400 MHz, $CDCl_3$) compound **2**

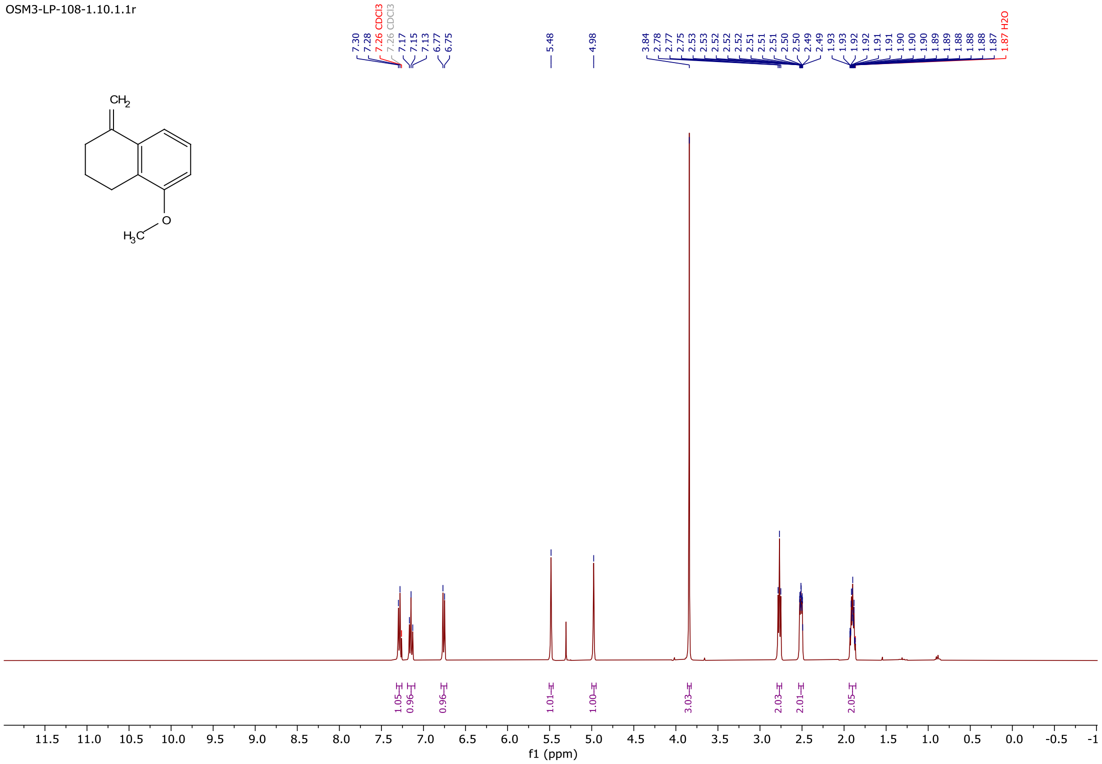


#### $^1$H NMR (400 MHz, $CDCl_3$) compound **3**

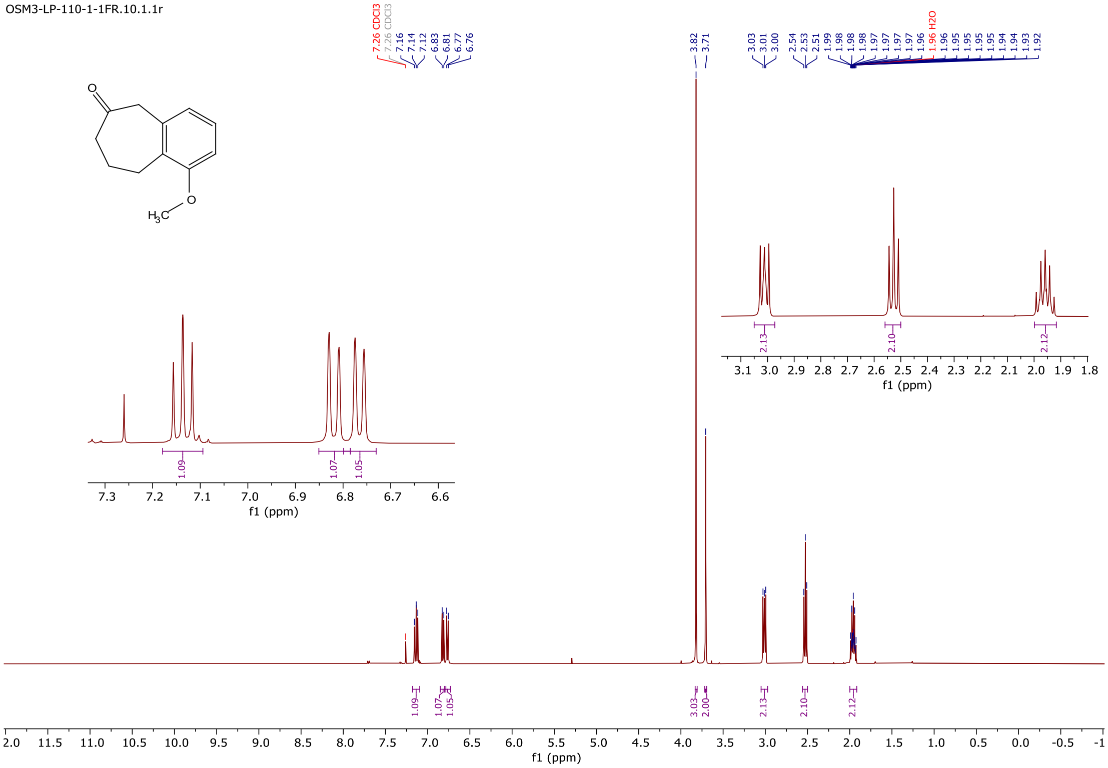

### $^{1}H$ NMR (400 MHz, $CDCl_3$) compound **4**

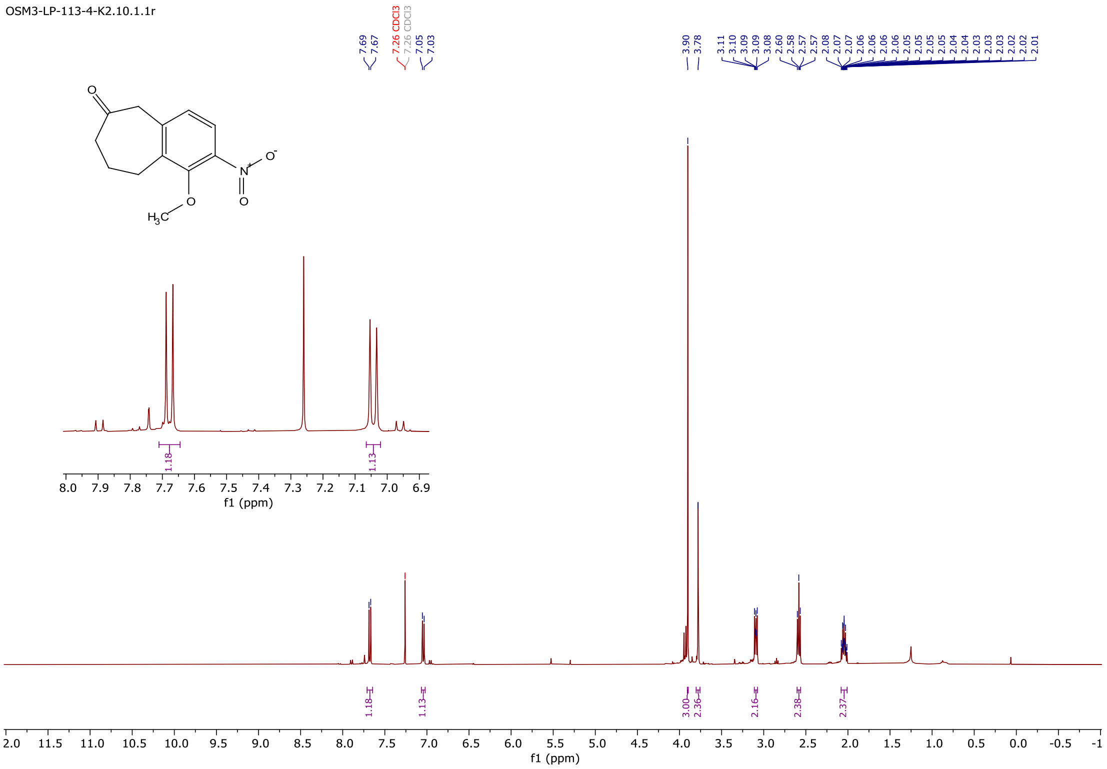


### $^{1}H$ NMR (400 MHz, $CDCl_3$) compound **6**

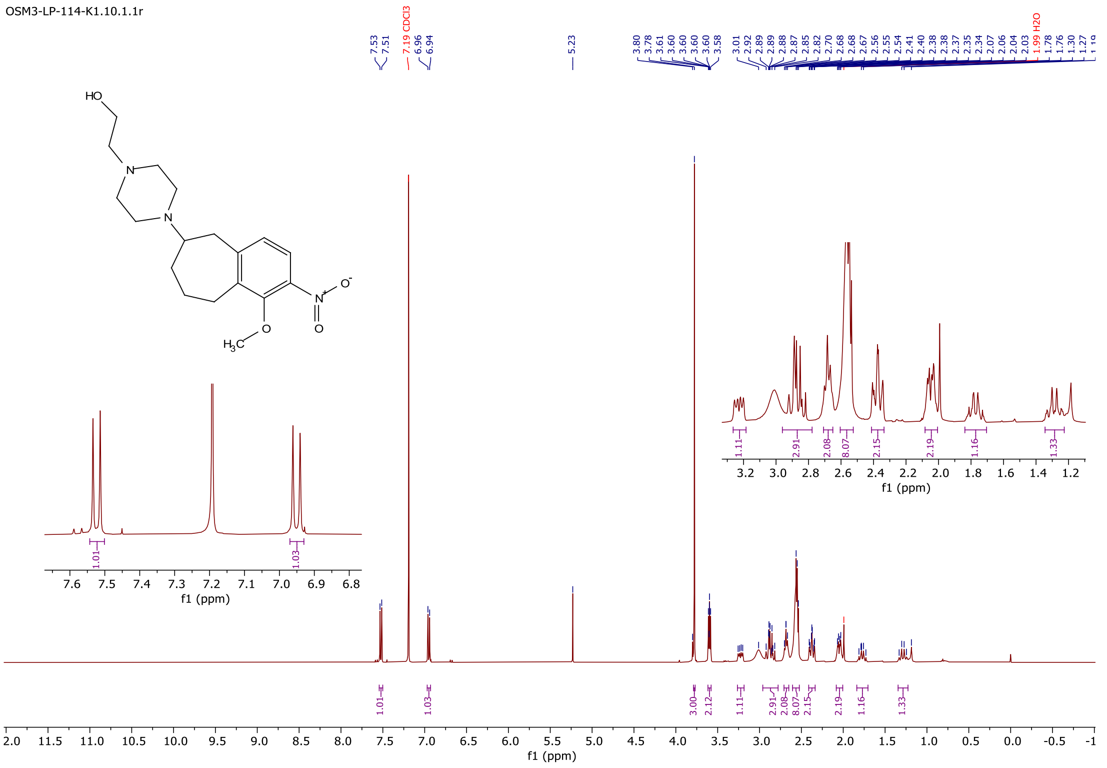

### $^{1}$H NMR (400 MHz, $CDCl_3$) compound **LP-115**

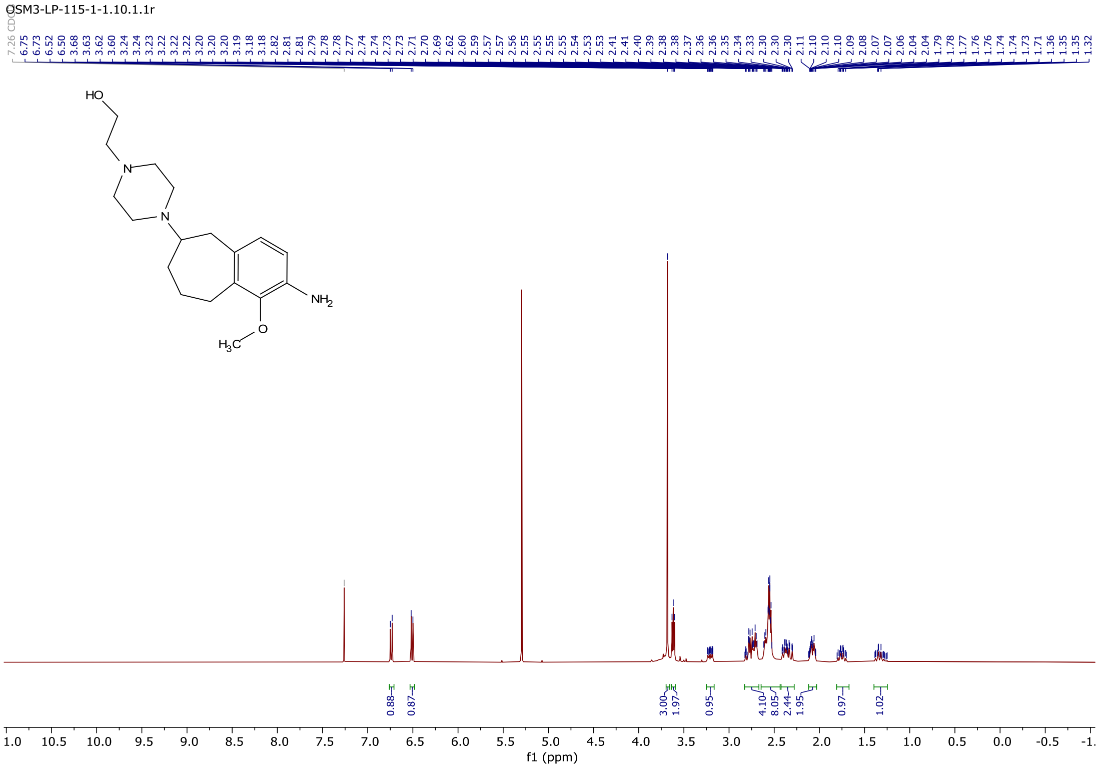


### $^{1}$H NMR (400 MHz, $CDCl_3$) compound **LP-117**

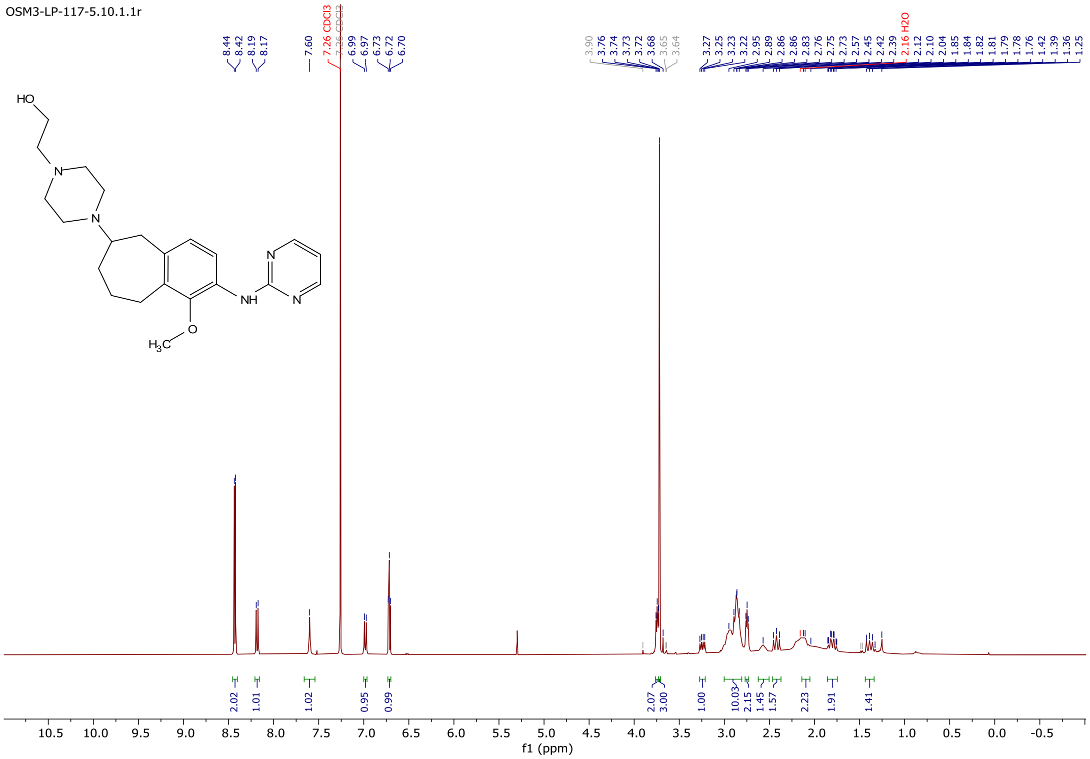

#### $^{13}C$ NMR (101 MHz, $CDCl_3$) compound **LP-117**

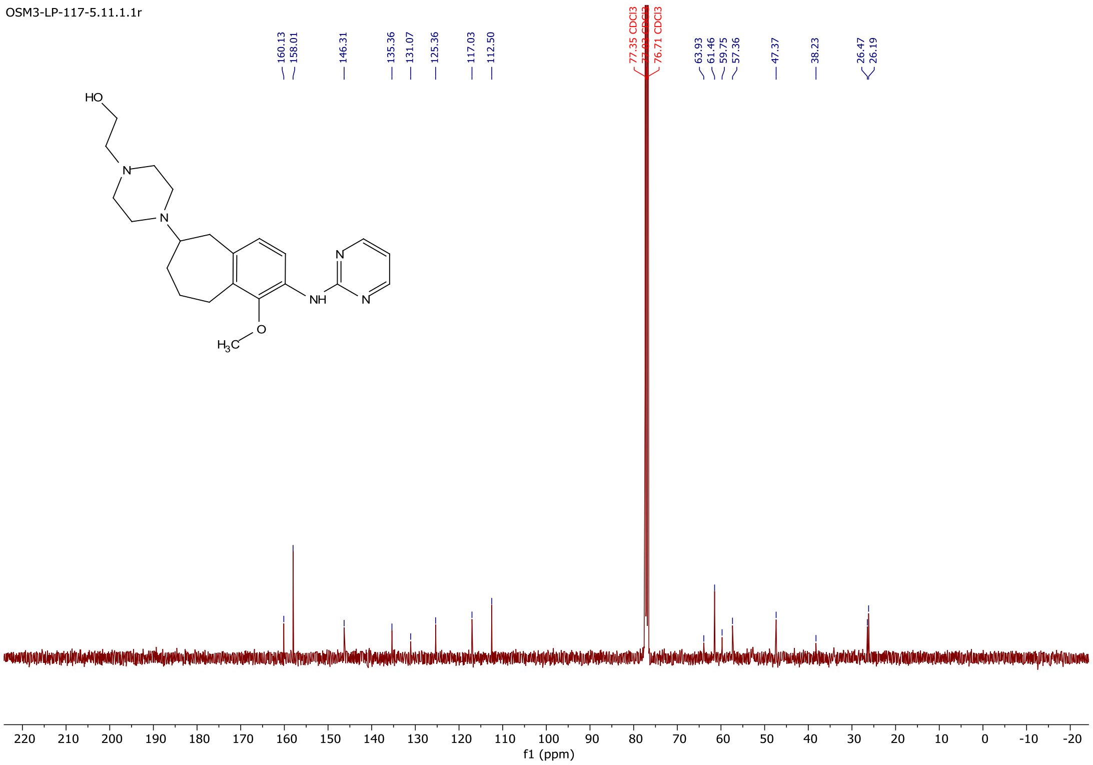


#### $^{1}H$ NMR (400 MHz, $CDCl_3$) compound **DGN-950**

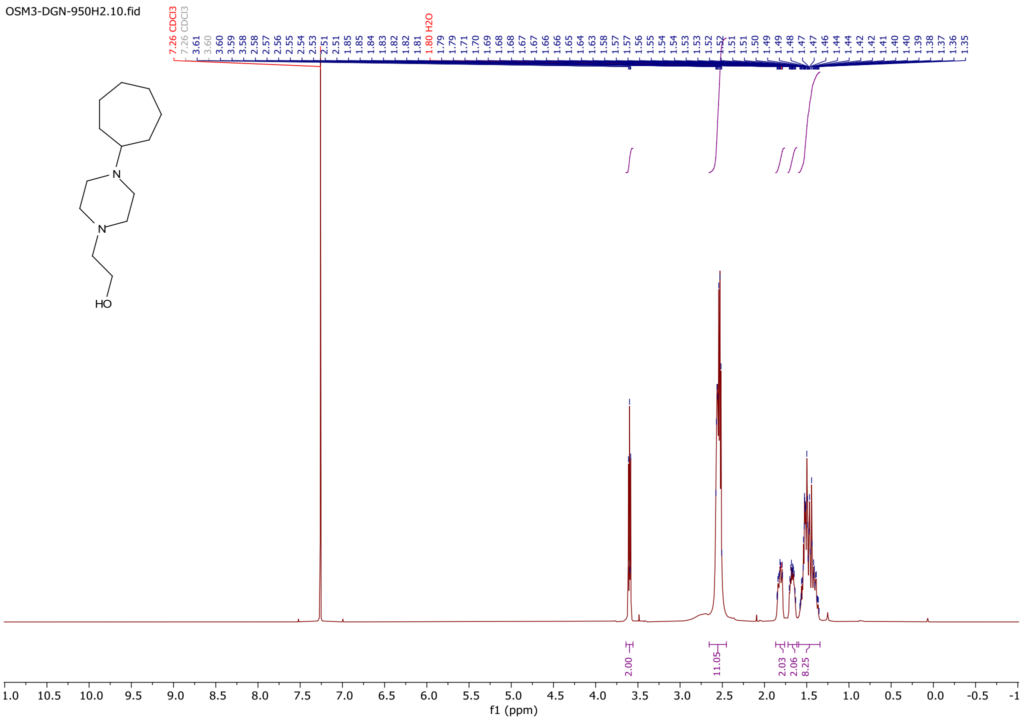

#### $^{13}C$ NMR (101 MHz, $CDCl_3$) compound **DGN-950**

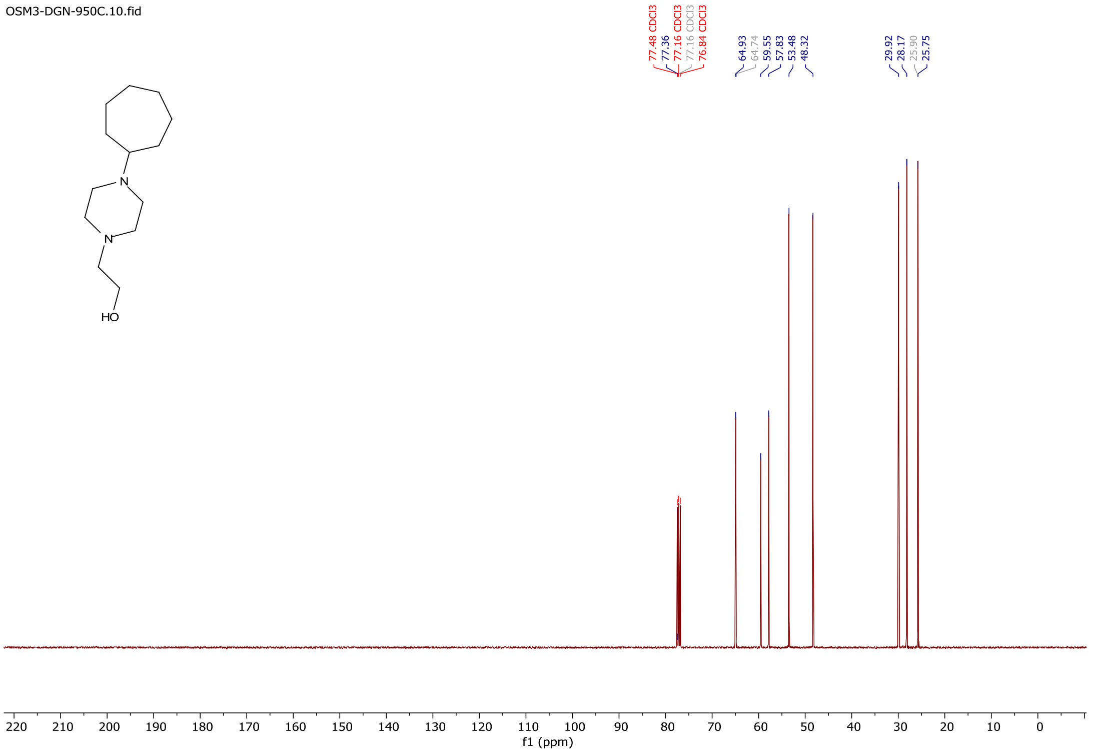


#### $^{1}H$ NMR (400 MHz, DMSO-$d_6$) compound **DGN-947**

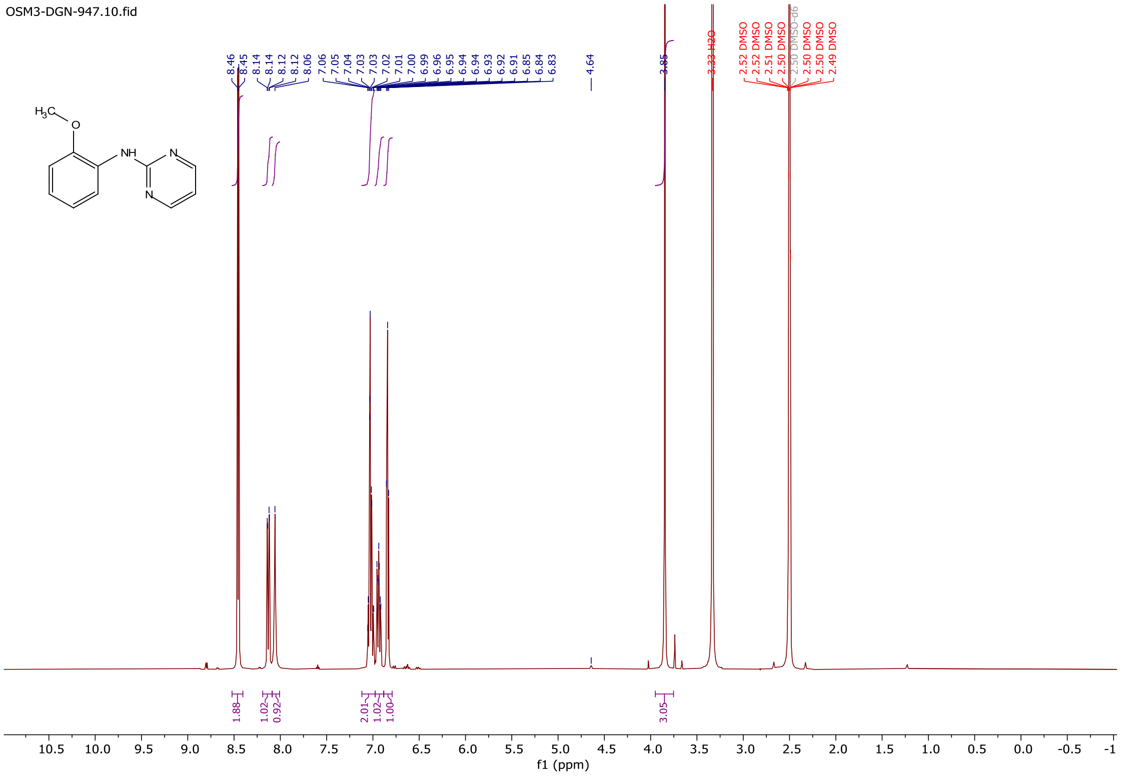

### $^{13}C$ NMR (101 MHz, DMSO-$d_6$) compound **DGN-947**

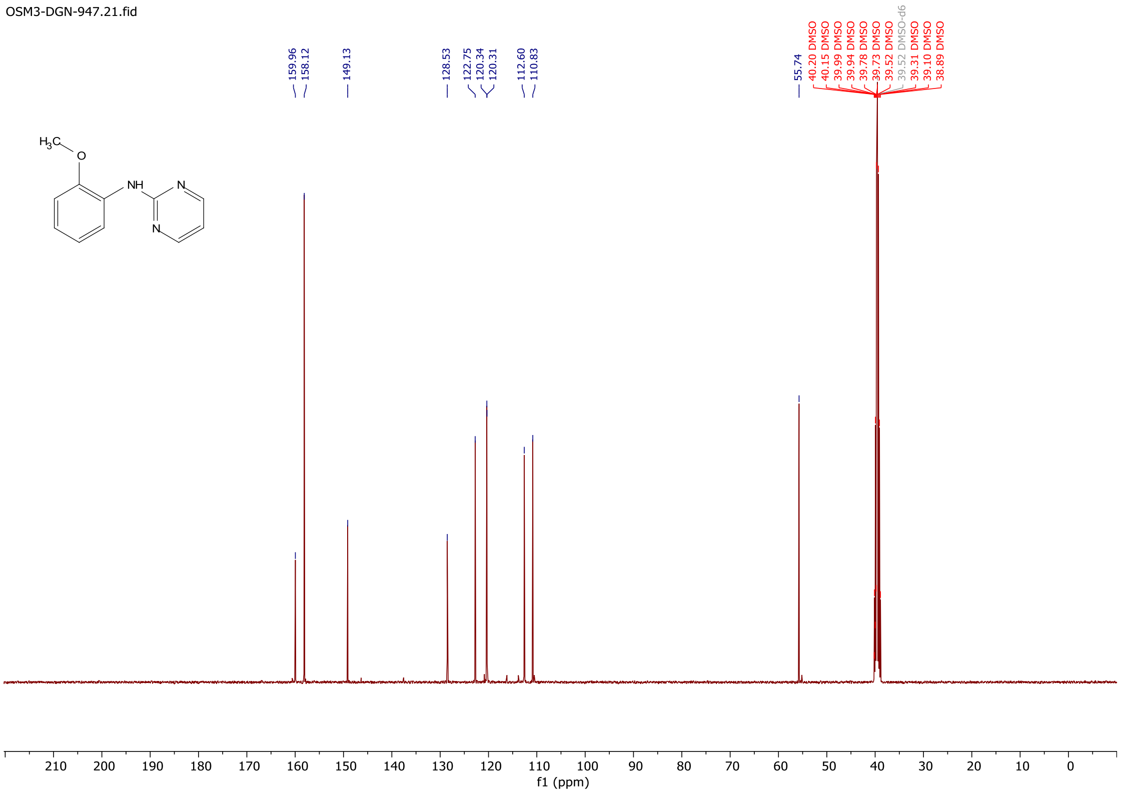
